# NTR 2.0: a rationally-engineered prodrug converting enzyme with substantially enhanced efficacy for targeted cell ablation

**DOI:** 10.1101/2020.05.22.111427

**Authors:** Abigail V. Sharrock, Timothy S. Mulligan, Kelsi R. Hall, Elsie M. Williams, David T. White, Liyun Zhang, Frazer Matthews, Saumya Nimmagadda, Selena Washington, Katherine Le, Danielle Meir-Levi, Olivia L. Cox, Meera T. Saxena, Anne L. Calof, Martha E. Lopez-Burks, Arthur D. Lander, Ding Ding, Hongkai Ji, David F. Ackerley, Jeff S. Mumm

## Abstract

Heterologously-expressed bacterial nitroreductase (NTR) enzymes sensitize eukaryotic cells to prodrugs such as metronidazole (MTZ), enabling selective cell ablation paradigms that have expanded studies of cell function and regeneration in vertebrate systems. However, first-generation NTRs require confoundingly toxic prodrug treatments (e.g. 10 mM MTZ) and some cell types have proven resistant. We used rational engineering and cross-species screening to develop a NTR variant, NTR 2.0, which exhibits ~100-fold improvement in MTZ-mediated cell-specific ablation efficacy. Toxicity tests in zebrafish showed no deleterious effects of prolonged MTZ treatments of ≤1 mM. NTR 2.0 therefore enables sustained cell loss paradigms and ablation of previously resistant cell types. These properties permit enhanced interrogations of cell function, extended challenges to the regenerative capacities of discrete stem cell niches, and enable modeling of chronic degenerative diseases. Accordingly, we have created a series of bipartite transgenic resources to facilitate dissemination of NTR 2.0 to the research community.

## INTRODUCTION

Bacterial nitroreductases (NTRs) are promiscuous enzymes capable of prodrug conversion via reduction of nitro substituents on aromatic rings^1–4^*. This generates genotoxic products that rapidly kill the host cell, a mechanism exploited by anti-cancer and antibiotic prodrugs^5^. When expressed heterologously, NTRs sensitize vertebrate cells to such prodrugs^6^. The canonical NTR, *Escherichia coli* NfsB (NfsB_Ec, “NTR 1.0”), has been widely tested in combination with the anti-cancer prodrug 5-(aziridin-1-yl)-2,4-dinitrobenzamide (CB1954) as an enzyme-prodrug therapy for killing tumors^1^. Transgenic expression of NTR 1.0 in combination with CB1954 was previously advanced as a targeted cell ablation strategy for interrogating cell function in vertebrates^7,8^. However, CB1954 produces cell-permeable cytotoxins that kill nearby non-targeted cells, i.e., “bystander” cell death^9^, compromising its use for selective cell ablation. In contrast, the prodrug metronidazole (MTZ) ablates NTR-expressing cells without discernible bystander effects^9^.

Importantly, fusion proteins between NTR and fluorescent reporters retain MTZ-inducible cell-specific ablation activity^10^. We therefore adapted the NTR/MTZ ablation system to zebrafish^11^ to expand studies of cellular regeneration^12^, reasoning that co-expression of NTR with reporters would enable visualization^13,14^ and quantification^14–17^ of MTZ-induced cell loss, and subsequent cell replacement, *in vivo*. In general, the NTR/MTZ system has proven useful for targeted cell ablation, and has had widespread uptake^12^. However, the low catalytic efficiency of NTR 1.0 requires high concentrations of MTZ (e.g., 10 mM) for effective cell ablation. This precludes sustained ablation paradigms, as MTZ exposures >24 h become increasingly toxic^18^. In addition, some cell types have proven resistant to NTR/MTZ-mediated ablation (e.g. macrophages and dopaminergic neurons)^19,20^. To overcome these limitations, we implemented a screening cascade to identify NTR variants exhibiting enhanced MTZ conversion activity.

Using targeted mutagenesis and high-throughput selection, the Searle group previously identified a triple mutant of NTR 1.0 exhibiting a 40-80-fold improvement in CB1954 conversion activity (NfsB_Ec T41Q/N71S/F124T, here “NTR 1.1”)^21^. This variant also enhanced MTZ-induced cell ablation in zebrafish, however, the improvement was only marginal (2-3 fold over NTR 1.0)^18,22^. In contrast, by leveraging a combination of rational engineering and cross-species testing, we identified a rationally engineered NfsB ortholog from the bacterial species *Vibrio vulnificus* (NfsB_Vv F70A/F108Y, “NTR 2.0”) which improves MTZ-meditated cell ablation efficiency ~100-fold. Additional data show that NTR 2.0 will expand the functionality of the NTR/MTZ system by allowing: 1) sustained interrogations of cell function, 2) effective ablation of “resistant” cell types, 3) prolonged cell loss, as novel tests of regenerative capacity, and 4) modeling of degenerative diseases caused by chronic cell loss. Accordingly, we have created a series of bipartite expression vectors and transgenic zebrafish lines co-expressing NTR 2.0 and fluorescent reporters as versatile new toolsets for the research community.

## RESULTS

### Rational improvement of metronidazole-activating NfsB_Ec-like nitroreductase variants

We previously compiled an *E. coli* gene library of 11 *nfsB* orthologs and used a DNA damage screen in *E. coli* host cells to monitor activation of SN33623, a PET imaging probe that shares a 5-nitroimidazole core structure with MTZ^23^. This same library was used here to evaluate ablation efficacy at higher SN33623 doses; relative growth of replicate *E. coli* cultures was assessed across a dilution series to establish EC_50_ values. Consistent with the previous DNA damage screen, the six most closely related orthologs of NTR 1.0 (‘NfsB_Ec-like’, >50% amino acid identity) were far less effective at activating SN33623 than the other five enzymes in the panel (**Fig. 1a,b**). MTZ sensitivity followed the same trend, with one notable exception: the *V. vulnificus* ortholog (NfsB_Vv), despite being NfsB_Ec-like, was one of the most active MTZ-converting enzymes (**Fig. 1c**).

**Fig. 1:**
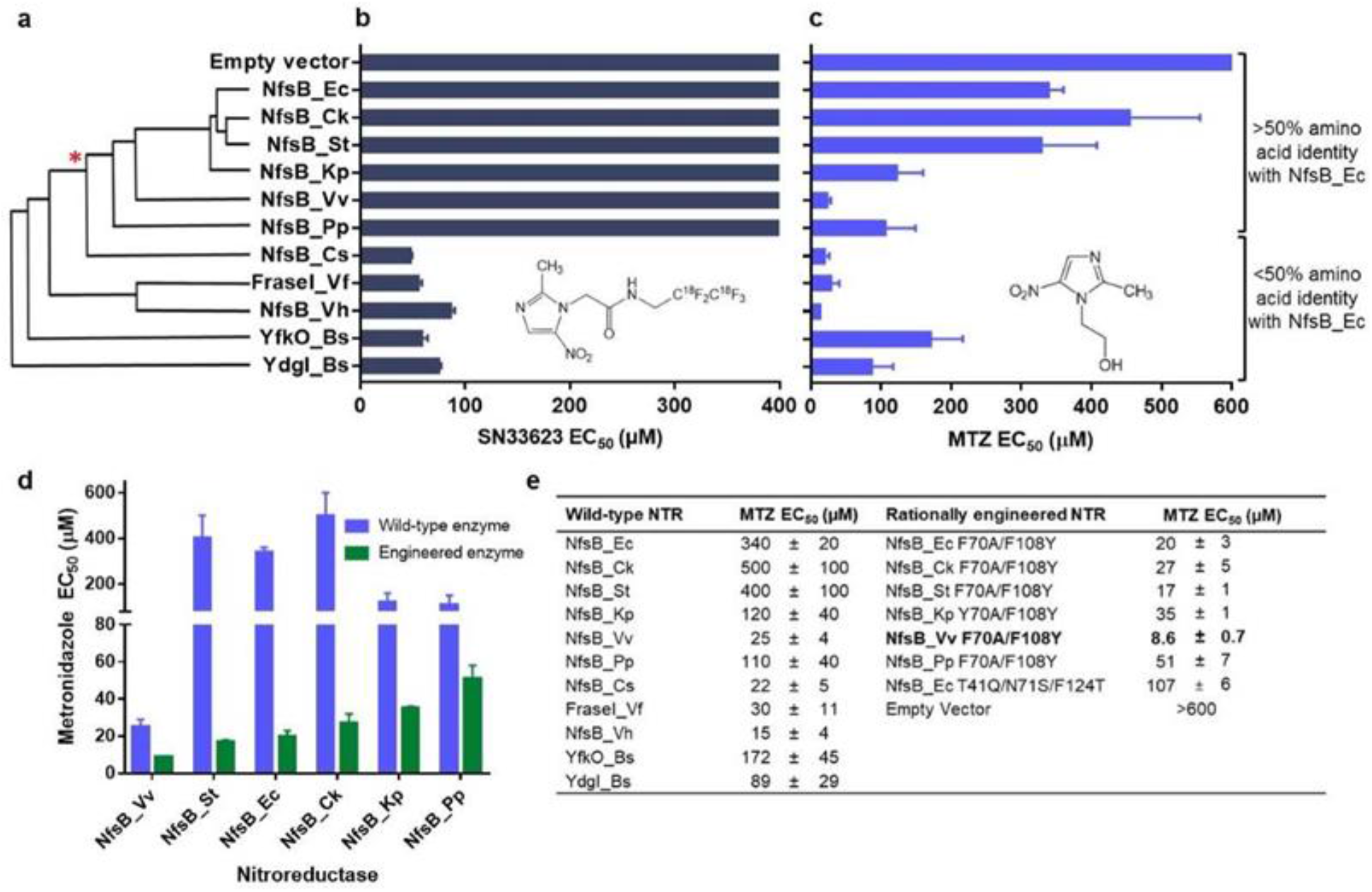
Rational engineering of NfsB-family nitroreductases for improved activation of metronidazole. **a,** Amino acid sequence identity cladogram of the eleven NfsB orthologs in our panel, grouped according to percent shared amino acid identity with NfsB_Ec. The asterisk (*) marks where other NTR variants diverge from NfsB_Ec-like enzymes (ClustalW2 tool with default settings; http://www.ebi.ac.uk/Tools/msa/clustalw2/). **b,c,** *E. coli* host sensitivity conferred by NfsB enzyme variants to the compounds SN33623 **(b)** or MTZ **(c)**; data without error bars indicate host cell sensitivity could not be observed within the tested concentration range. *Insets*: chemical structures of SN33623 and MTZ. **d,** *E. coli* host sensitivity to MTZ conferred by wild-type or rationally engineered NfsB-like enzymes (i.e., with F70A/F108Y substitutions). **e,** Summary of MTZ EC_50_ values for *E. coli* strains expressing wild-type or rationally engineered NTRs. Data are the averages of at least three biological replicates ± SD.

This was a promising finding, as loss-of-activity and gain-of-activity rational mutagenesis previously demonstrated that the residues F70 and F108 impair 5-nitroimidazole activity in NfsB_Ec^23^, and these residues are highly conserved in the NfsB_Ec-like enzymes (**Supplementary Fig. S1**). We hypothesized that introducing F70A/F108Y substitutions into NfsB_Vv and the other NfsB_Ec-like enzymes (Y70A/F108Y for NfsB_Pp) would enhance MTZ activity, as was previously found for NfsB_Ec^23^. This proved true in all cases, with host cells expressing the substituted variants being substantially more sensitive to MTZ. The *E. coli* strain expressing NfsB_Vv F70A/F108Y was the most sensitive to MTZ, having an EC_50_ 40-fold lower than NfsB_Ec (NTR 1.0) and 12-fold lower than NfsB_Ec T41Q/N71S/F124T (NTR1.1) expressing strains (**Fig. 1d,e**).

To determine whether *E. coli* EC_50_ data accurately reflected improved enzymatic reduction of MTZ, the native and F70A/F108Y substituted NfsB_Vv variants were purified as His_6_-tagged proteins using nickel affinity chromatography, and MTZ conversion activities were compared to the benchmark NTR 1.0 and NTR 1.1 enzymes. Relative Michaelis-Menten kinetic parameters were consistent with *E. coli* sensitivity data; NfsB_Vv F70A/F108Y exhibited the lowest *K*_*M*_ and highest catalytic efficiency (*k*_*cat*_/*K*_*M*_), followed by NfsB_Vv, NTR 1.1, and NTR 1.0 respectively (**Supplementary Fig. S2**; **Supplementary Table S1**).

### Cell ablation activity of NfsB_Vv F70A/F108Y *in vitro*: mammalian cells

To assess relative MTZ conversion activities (and expression tolerance) of lead NTR variants in vertebrate cells, NfsB_Vv F70A/F108Y, NfsB_Vv, NTR 1.0 and NTR 1.1 were stably transfected into human cells (HEK-293). MTZ sensitivity was evaluated using a MTS viability assay^24^ following a 48 h MTZ dose-response challenge (5 mM to 1 μM, 2-fold dilution series). Cells expressing NTR 1.1 lost expression over time, reflected in the comparatively high MTZ EC_50_ value (2300 μM). In contrast, NfsB_Vv F70A/F108Y, NfsB_Vv and NTR 1.0 were stably expressed and relative MTZ sensitivities (3, 9 and 1700 μM respectively) were concordant with the *E. coli* EC_50_ data. Equivalent MTZ dose-response assays, using the MTT viability assay^25^, were performed on Chinese hamster ovary (CHO-K1) cell lines expressing either NTR 1.1 or NfsB_Vv F70A/F108Y. The resulting EC_50_ values of 690 μM and 4 μM, respectively, suggest the improved MTZ activity exhibited by NfsB_Vv F70A/F108Y is retained in heterologous mammalian expression systems (**Supplementary Fig. S3**; **Supplementary Table S2**).

### Ablation specificity of NfsB_Vv F70A/F108Y *in vitro*: mammalian cells

A key advantage of MTZ is the cell-specific nature of its cytotoxic metabolite, allowing selective ablation of NTR-expressing cells without harming surrounding cells. This targeted ablation activity enables precise elimination of cellular subtypes and subsequent delineation of roles during developmental, regenerative, and other biological processes of interest. To test NfsB_Vv F70A/F108Y mediated ablation specificity, HEK-293 cell lines were generated that stably expressed either green fluorescent protein (GFP) alone or NfsB_Vv F70A/F108Y and mCherry. When single cultures were challenged with 6 μM MTZ or 0.01% (v/v) DMSO (control) for 48 h, cells expressing NfsB_Vv F70A/F108Y and mCherry (**Fig. 2a,a’**) were killed, whereas cells expressing GFP remained healthy (**Fig. 2b,b’**). Co-cultures undergoing the same challenge directly demonstrated selective ablation of NfsB_Vv F70A/F108Y-expressing cells (**Fig. 2c,c’**). At this point, having confirmed that targeted cell ablation activity was enhanced in both bacterial and vertebrate cell lines, we dubbed the engineered NfsB_Vv F70A/F108Y variant “NTR 2.0”.

**Fig. 2:**
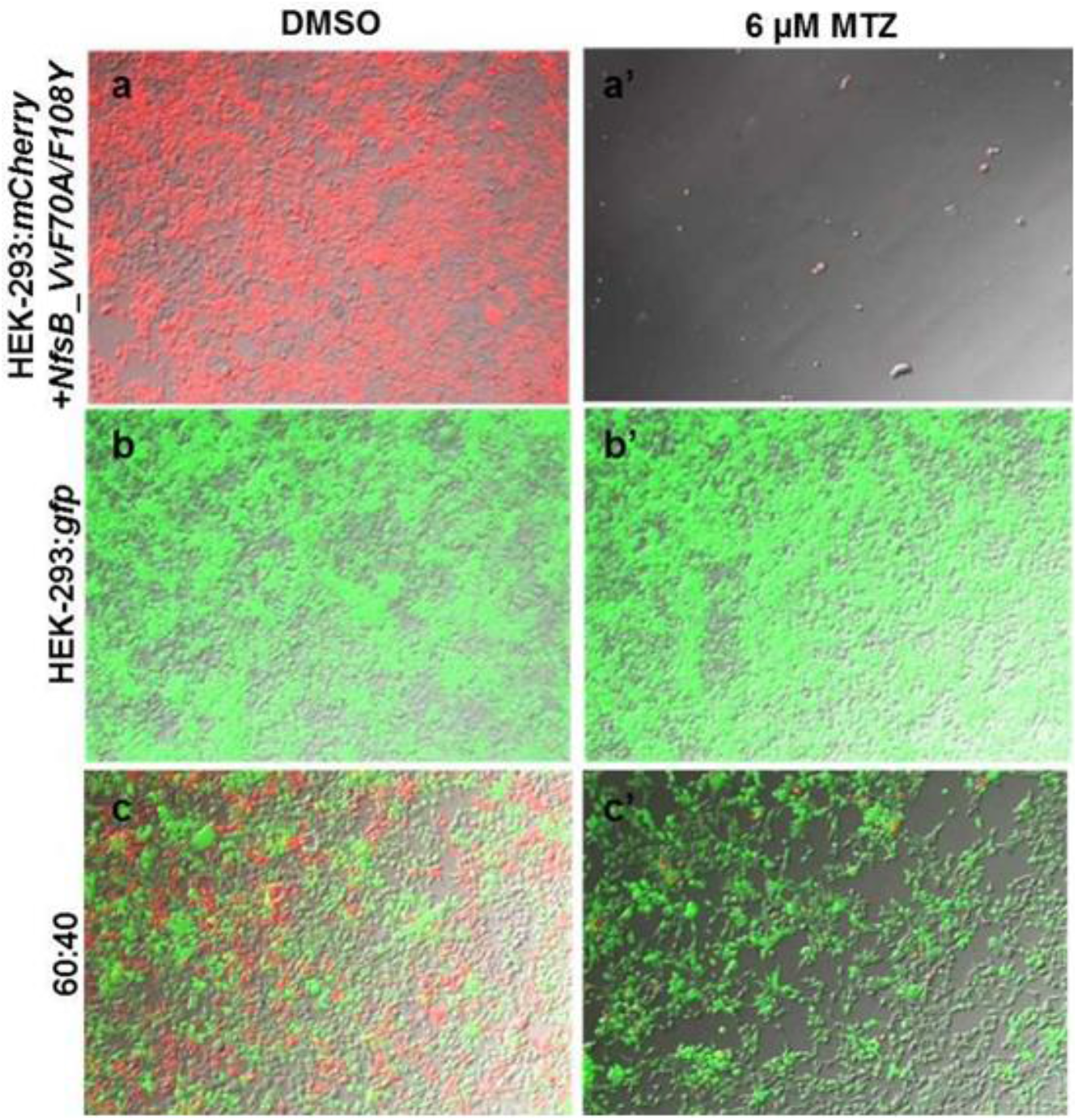
NfsB_Vv F70A/F108Y conversion of metronidazole retains targeted ablation activity. **a-c,** HEK-293 cell lines expressing GFP (HEK-293:*gfp*) **(a)** or NfsB_Vv F70A/F108Y and mCherry (HEK-293:*mCherry+nfsB_Vv F70AF108Y*) **(b)** were grown in isolation **(a,b)** or in a 60:40 co-culture **(c)**. Cells were treated with 0.01% DMSO control **(a-c)** or 6 μM MTZ **(a’-c’)** for 48 h. Cell viability was assessed qualitatively by fluorescence microscopy.

### Cell ablation activity of NfsB_Vv F70A/F108Y (NTR 2.0) *in vivo*: zebrafish assays

To determine if NTR 2.0 increases MTZ-induced ablation efficacy *in vivo*, we generated a UAS reporter/effector transgenic zebrafish line co-expressing yellow fluorescent protein (YFP) and NTR 2.0 in a Gal4-dependent manner, *Tg(5xUAS:GAP-tagYFP-P2A-nfsB_Vv F70A/F108Y)jh513* (*UAS:YFP-NTR2.0*; see Methods for full list of transgenic zebrafish lines used here). This line was crossed with a neuronally-restricted Gal4 driver line^26^, *Et(2xNRSE-fos:KalTA4)gmc617* (KalTA4 is a zebrafish optimized Gal4/VP16 fusion^27^), to establish double transgenic larvae co-expressing YFP and NTR 2.0 (*NRSE:KalTA4;UAS:YFP-NTR2.0)* throughout the nervous system (**Supplementary Fig. S4**).

To test NTR 2.0 for enhanced cell ablation efficacy *in vivo*, 5 day post-fertilization (dpf) double-transgenic *NRSE:KalTA4;UAS:YFP-NTR2.0* larvae were exposed to MTZ for 24 h across a 2-fold dilution series from 200 to 12.5 μM. Using an established plate reader assay^15,16^, YFP levels were quantified in individual fish at 7 dpf. YFP was undetectable in transgenic fish exposed to 200 μM MTZ, i.e., reduced to non-transgenic control levels (**Fig. 3a**). Dose-dependent effects were evident between 25 and 200 μM MTZ with a calculated EC_50_ of 39 μM. This represents a ~100-fold improvement over NTR 1.0 activity and ~50-fold improvement over NTR 1.1^18^. Extending MTZ treatments to 48 h resulted in a more pronounced dose-dependent effect and an EC_50_ of 16 μM (**Fig. 3b**). To determine if NTR 2.0 enabled ablation upon shorter exposures to higher MTZ concentrations, 5 dpf transgenic larvae were incubated with 0.5, 1 or 10 mM MTZ for 2 h or 24 h (control). A 2 h incubation with 1 mM or 10 mM MTZ resulted in near identical levels of ablation relative to 24 h controls (**Fig. 3c**; **Supplementary Table S3**).

**Fig. 3:**
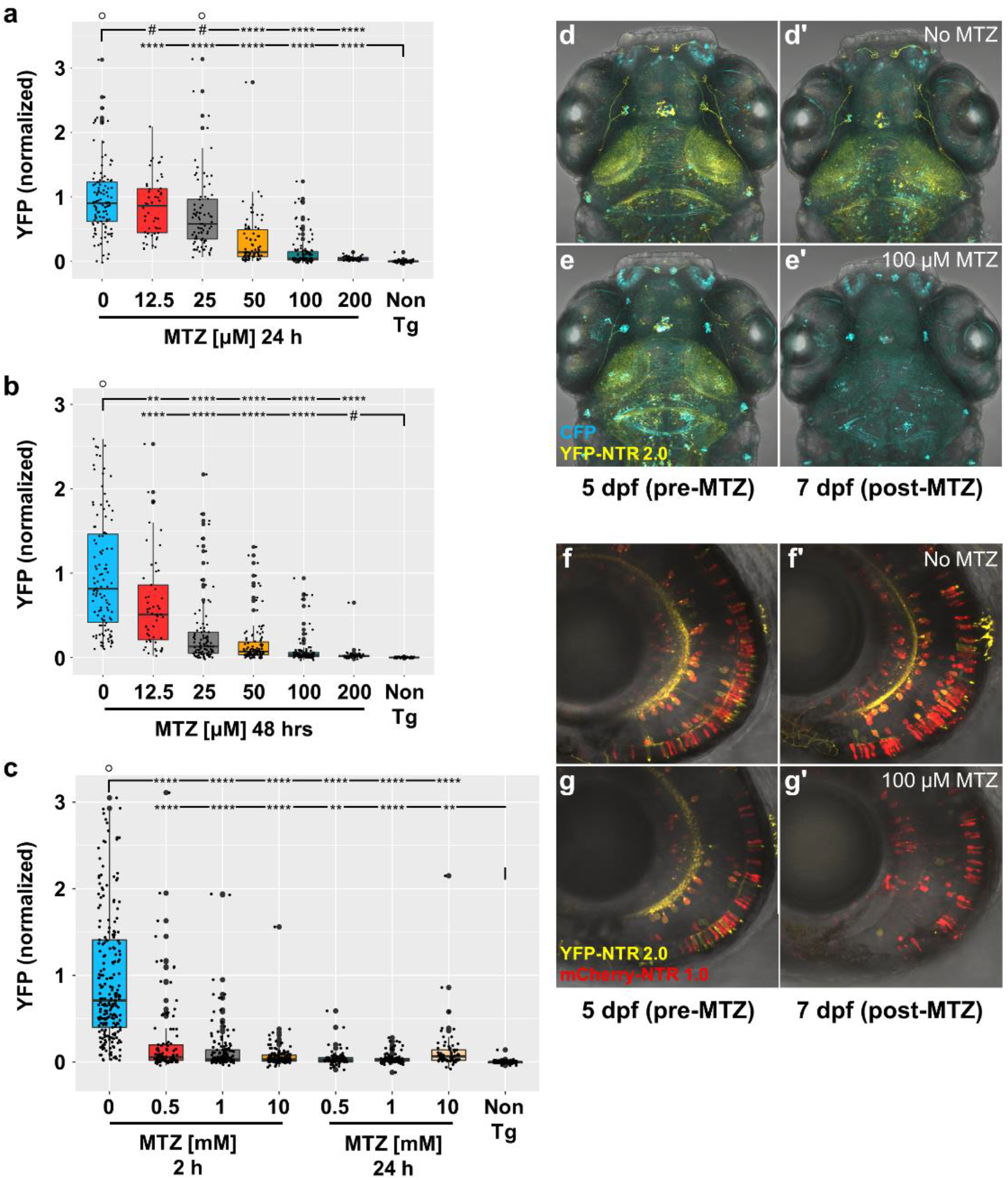
NTR 2.0 enhances cell-specific ablation efficacy *in vivo*. **a-c**, Dose-response tests of MTZ ablation efficacy. Double transgenic zebrafish larvae co-expressing YFP and NTR 2.0 were exposed to the indicated concentrations of MTZ for 24 h **(a)**, 48 h **(b)**, or 2 h **(c)**. Controls included transgenic larvae not exposed to MTZ (0) and non-transgenic larvae (Non-Tg), providing upper and lower YFP expression level bounds, respectively. YFP levels were quantified using an established plate reader assay^15,16^. Statistical comparisons to two controls, non-ablated (0 mM MTZ) and age-matched non-transgenic (Non-Tg) larvae, were used to calculate absolute effects sizes, 95% confidence intervals, and Bonferroni-corrected *p’*-values (provided in Supplementary Table S3; ^#^*p*>0.05, \**p*<0.05, ** *p*<0.01, *** *p*<0.001, **** p<0.0001; ○ = data points beyond the bounds of the graph). **d,e,** Ablation specificity test of NTR 2.0 *in vivo*. Triple transgenic zebrafish larvae expressing YFP and NTR 2.0 (yellow cells) and/or CFP (blue cells) were exposed to 100 μM MTZ from 5-6 dpf. Time series confocal imaging of dorsal brain regions showed overlapping (YFP/CFP cells) and non-overlapping (YFP or CFP only) expression prior to MTZ exposure at 5 dpf **(d,e)**. Control larvae (No MTZ) imaged at 7 dpf showed retention of both YFP-NTR 2.0 and CFP expressing cells **(d’)**. MTZ-treated larvae showed loss of YFP-NTR 2.0 expressing cells but sustained expression of CFP-only expressing cells (**e’**). **f,g,** Ablation efficacy comparison of NTR 1.0 and NTR 2.0. Triple transgenic zebrafish larvae expressing YFP and NTR 2.0 (yellow cells) and/or mCherry and NTR 1.0 (red cells) were exposed to 100 μM MTZ for 48 h from 5-7 dpf. Time series confocal imaging of retinal neurons showed overlapping (YFP/mCherry cells) and non-overlapping (YFP or mCherry only) expression prior to MTZ exposure at 5 dpf **(f,g)**. Control larvae (No MTZ) imaged at 7 dpf showed retention of both YFP-NTR 2.0 and mCherry-NTR 1.0 expressing cells **(f’)**. MTZ-treated larvae showed loss of YFP-NTR 2.0 expressing cells but sustained expression of mCherry-NTR 1.0 expressing cells **(g’)**.

### Ablation specificity of NTR 2.0-converted MTZ *in vivo*: zebrafish

To test NTR 2.0/MTZ-induced ablation specificity *in vivo*, we generated triple transgenic fish combining the *NRSE:KalTA4*;*UAS:YFP-NTR2.0* line (**Supplementary Fig. S4**) with a second UAS reporter expressing CFP, *Tg(5xUAS:GAP-ECFP,he:GAP-ECFP)gmc1913* (*UAS:CFP*). Due to mosaicism, the two UAS reporters are expressed in overlapping and non-overlapping subsets; non-targeted cells expressing CFP only and targeted cells expressing YFP-NTR 2.0 with or without CFP (**Fig. 3d,e**). In the absence of MTZ, mosaic expression of both reporters is maintained from 5 to 7 dpf (**Fig. 3d,d’**). Fish exposed to 100 μM MTZ (24 h), however, exhibited selective loss of cells expressing YFP and NTR 2.0 while cells expressing CFP alone remained (**Fig. 3e,e’**). This result confirms that NTR 2.0 confers selective cell ablation activity *in vivo*.

To directly compare ablation efficacies of NTR 1.0 and NTR 2.0 *in vivo*, we generated triple transgenic fish combining *NRSE:KalTA4*;*UAS:YFP-NTR2.0* lines with a second UAS reporter expressing a NTR 1.0-mCherry fusion protein, *Tg(14xUAS: nfsB_Ec-mCherry)c264*^28^ (*UAS:NTR1.0-Cherry*). Mosaicism again led to overlapping and non-overlapping expression of UAS transgenes (**Fig. 3f,g**). In the absence of MTZ, mosaic expression of both reporters is maintained from 5 to 7 dpf (**Fig. 3f,f’**). In contrast, YFP and NTR 2.0 expressing cells were selectively lost in larvae exposed to 100 μM MTZ for 24 h while cells expressing Cherry and NTR 1.0 alone remained (**Fig. 3g,g’**). This result further confirms the superior cell ablation efficacy of NTR 2.0.

To further test for enhanced ablation efficacy, we compared two additional transgenic lines expressing NTR 1.0 or NTR 2.0 in retinal rod photoreceptor cells, *Tg(rho:YFP-nfsB_Ec)gmc500* (*rho:YFP-NTR1.0*) and *Tg(rho:GAP-YFP-2A-nfsB_Vv F70A/F108Y)jh405* (*rho:YFP-NTR2.0*), respectively. In dose-response tests (five-fold dilution series, from 5 mM - 0.32 μM, 48 h treatment), the NTR 1.0-expressing line produced an EC_50_ of 540 μM MTZ (**Fig. 4a**), in keeping with our prior observation that rod photoreceptors can be ablated with relatively lower MTZ concentrations (2.5 mM)^17^. The NTR 2.0-expressing line, however, had an EC_50_ of 3 μM MTZ, equating to ~180-fold improvement in ablation efficacy (**Fig. 4b**; **Supplementary Table S4**).

**Fig. 4:**
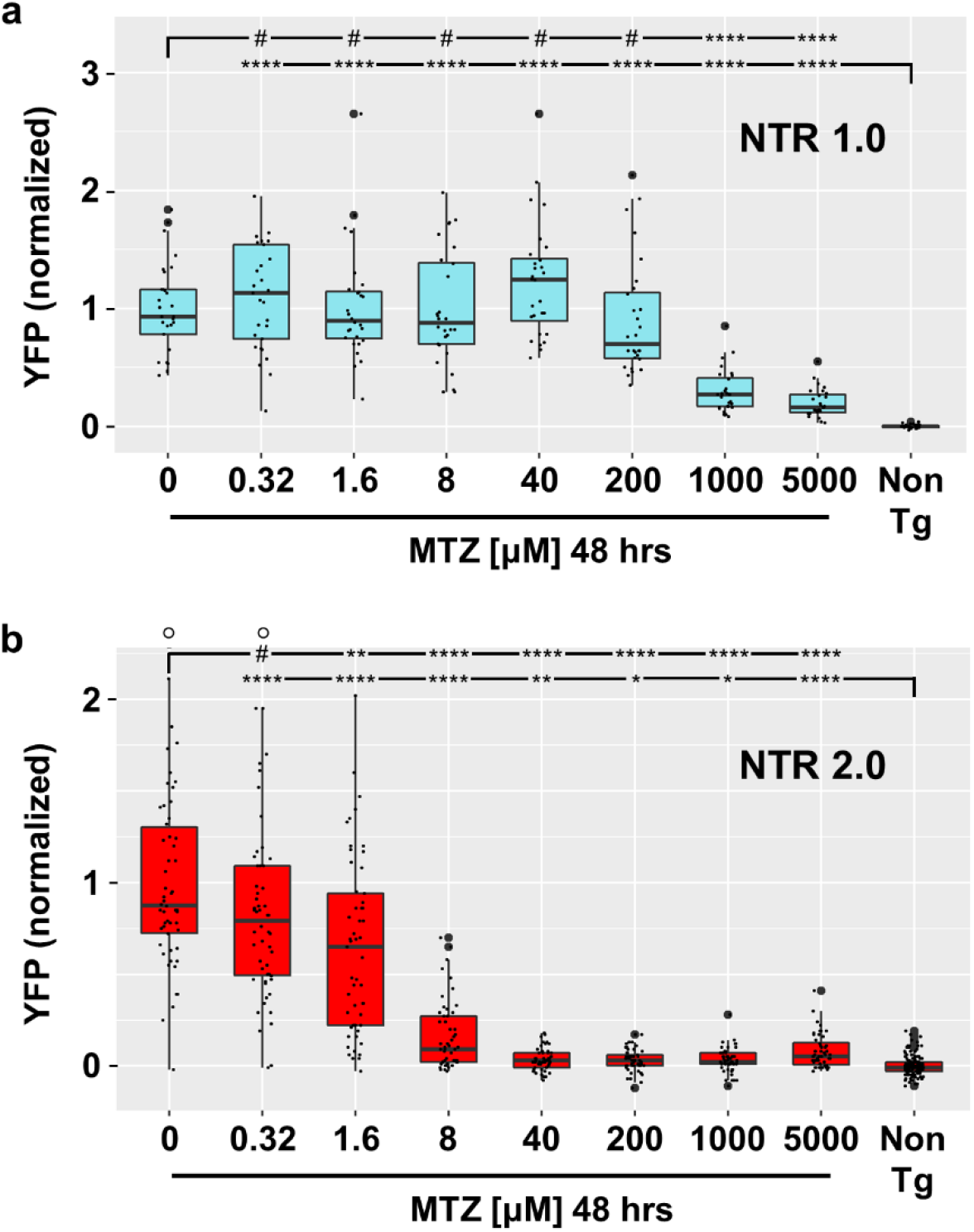
Dose-response of rod cell ablation effic – NTR 1.0 versus NTR 2.0. **a,b,** Transgenic zebrafish larvae co-expressing either NTR 1.0 and YFP **(a)** or NTR 2.0 and YFP **(b)** were treated with MTZ across a 5-fold dilution series (5 mM – 320 nM) from 5-7 dpf. YFP levels were quantified using an established plate reader assay. Statistical comparisons to two controls, non-ablated (0 mM MTZ) and age-matched non-transgenic (Non-Tg) larvae, were used to calculate absolute effect sizes, 95% confidence intervals, and Bonferroni-corrected *p’*-values (provided in Supplementary Table S4; ^#^*p*>0.05, \**p*<0.05, ** *p*<0.01, *** *p*<0.001, **** p<0.0001; ○ = data points beyond the bounds of the graph).

### Prolonged low dose MTZ exposures in zebrafish

A key disadvantage of NTR 1.0 is the near-toxic concentrations of MTZ needed to ablate cells. In addition to potential off-target effects^29^, near-toxic doses preclude sustained ablation paradigms^18^. To determine whether zebrafish can tolerate long-term low dose MTZ exposures, 15 dpf larvae were exposed to 10, 1, 0.1 or 0 mM MTZ for 36 days. As observed previously^18^, fish maintained in 10 mM MTZ exhibited high rates of lethality with only 6.1% alive after twelve days and none surviving beyond day 34 (**Fig. 5a**). In contrast, for all other conditions fish survival was equivalent, with ≥92% alive at day 36 (1 mM = 94%, 0.1 mM = 92%, 0 mM = 94%; **Fig. 5a**).

**Fig. 5:**
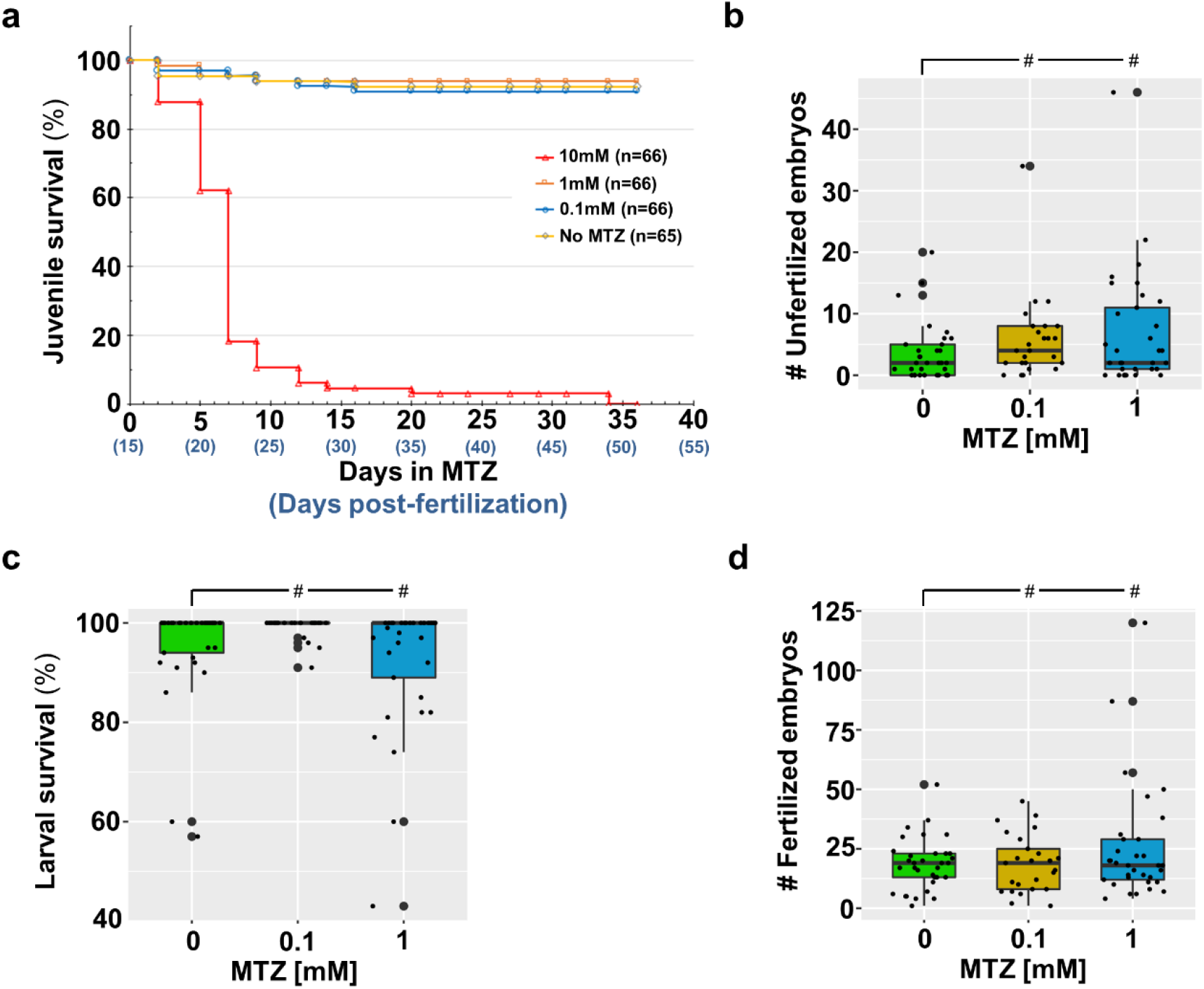
Effects of prolonged metronidazole incubation on juvenile and adult zebrafish. **a,** Survival of zebrafish incubated with the indicated concentration of MTZ for 36 days, 15-51 dpf. Log-rank (Mantel-Cox) tests and Gehan-Breslow-Wilcoxon tests showed no statistically significant differences between the 0 mM MTZ control and 0.1 or 1 mM MTZ conditions, however, comparisons to the 10 mM condition produced chi-squares of 123 and 97, respectively, and a Bonferroni corrected *p’*-value of <0.0001 for both tests. **b-d,** At 3 months of age, fertility **(b)**, fecundity **(c)** and survival rates of offspring **(d)** were tested through pairwise in-crosses for each group. Data was subjected to ANOVA followed by Dunnett’s multiple comparisons test, depending on the outcome of Levene’s test. Comparisons to the 0 mM MTZ control group were used to calculate absolute effect sizes, 95% confidence intervals and *p-* values (provided in supplementary Table S5; ^#^*p*>0.05).

Long-term MTZ exposure has been implicated in rodent male infertility^30^ and associated (rarely) with neurotoxic effects in patients^31^. Therefore, we assessed fecundity, fertility, and offspring survival rates of fish exposed to 1, 0.1, or 0 mM MTZ once they reached sexual maturity (~3 months of age). No deleterious effects on clutch size, fertilization rates, or offspring survival rates were observed in MTZ-treated fish compared to untreated controls (**Fig. 5b-d**; **Supplementary Table S5**).

### NTR 2.0 enables ablation of NTR/MTZ resistant cell types

Another limitation of first-generation NTR resources is that certain cell types have proven difficult to ablate. For example, macrophages co-expressing YFP and NTR 1.1, *Tg(mpeg1.1:EYFP-nfsB_Ec T41Q/N71S/F124T)w202* (*mpeg:YFP-NTR1.1*), are resistant to 10 mM (24 h) doses of MTZ^14^, and effective ablation requires multi-day exposures to 2.5 mM MTZ in adult fish^20^. To determine if NTR 2.0 enables ablation of “resistant” cell types, double transgenic fish co-expressing YFP and NTR 2.0 in macrophages were generated by crossing a macrophage-specific KalTA4 driver, *Tg(mpeg1.1:Gal4-VP16)gl24*, with *UAS:YFP-NTR2.0* (*mpeg:Gal4;UAS:YFP-NTR2.0*). To assess relative macrophage ablation efficacy, *mpeg:YFP-NTR1.1* and *mpeg:Gal4;UAS:YFP-NTR2.0* larvae were exposed to 10, 1, 0.1, or 0 mM MTZ from 5 to 7 dpf. Confocal time series images show incomplete loss of NTR 1.1-expressing cells treated with 10 mM MTZ (**Fig. 6a**) but near complete ablation of cells expressing NTR 2.0 treated with 100 μM MTZ (**Fig. 6b**). Imaris-based volumetric quantification of YFP levels in confocal image stacks^14^ showed differential ablation efficacy for NTR 1.1 and NTR 2.0 across all MTZ treatment conditions (**Fig. 6b,d**). Lower doses proved entirely ineffective for ablating macrophages expressing NTR 1.1, and 10 mM MTZ resulted in only a 29% reduction (**Fig. 6b**; **Supplementary Table S6**). Conversely, more pronounced levels of macrophage ablation (64-73% reductions) were produced across all three doses of MTZ for cells expressing NTR 2.0 (**Fig. 6d**; **Supplementary Table S6**).

**Fig. 6:**
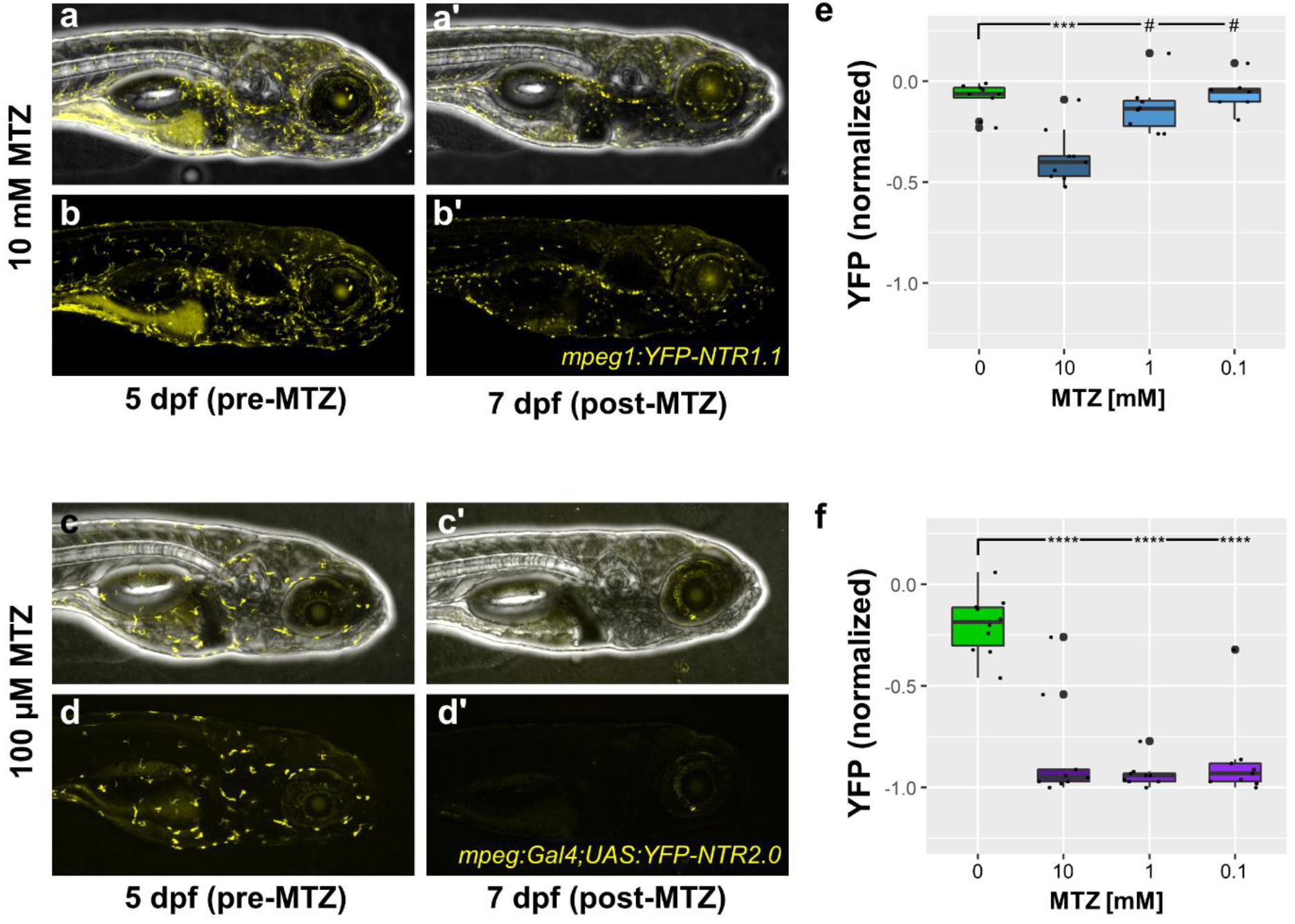
NTR 2.0 enables ablation of “resistant” cell types. **a,d,** Transgenic larvae expressing a YFP-NTR 1.1 fusion protein **(a,b)** or co-expressing YFP and NTR 2.0 **(c,d)** in macrophages/microglia were treated with 0, 0.1, 1, or 10 mM MTZ from 5-7 dpf. Intravital time series imaging was performed on pre- **(a-d)** and post-MTZ **(a’-d’)** treatment larvae at 5dpf and 7 dpf, respectively, and z-stack images were acquired with an Olympus FV1000 confocal microscope. **e,f,** Quantification of macrophage/microglia numbers in pre- and post-treatment images of larvae expressing either NTR 1.1 **(a,b series)** or NTR 2.0 **(c,d series)** was performed on 3D-rendered z-stacks (ImageJ v1.52h; NIH) using IMARIS v9.2 (Bitplane). The percent change in cell numbers was calculated by normalizing day 7 to day 5 image values per each fish (n=7-10 per condition). Note: the slight reduction of macrophage/microglia numbers in the untreated condition at day 7 is likely due to developmental changes. Statistical comparisons to the 0 mM MTZ control were used to calculate absolute effects sizes, 95% confidence intervals, and Bonferroni-corrected *p’*-values (provided in Supplementary Table S6; ^#^*p*>0.05, *** *p*<0.001, **** p<0.0001).

### NTR 2.0 does not improve nifurpirinol ablation efficacy

In addition to MTZ, NTR enzymes can convert other prodrug substrates. For instance, a recent study demonstrated robust NTR 1.1-mediated cell ablation following a 24 h exposure to 2.5 μM nifurpirinol (NFP)^32^. To determine if NTR 2.0 exhibits improved NFP conversion activity, double transgenic larvae co-expressing YFP and NTR 2.0 (*NRSE:KalTA4;UAS:YFP-NTR2.0*) were exposed to a 1:10 dilution series of nifurpirinol (2.5 μM - 2.5 nM) from 5 to 6 dpf, and YFP quantified at 7 dpf. Contrary to the ~100-fold improvement observed with MTZ, no appreciable enhancement of cell ablation efficacy was observed with nifurpirinol (**Supplementary Fig. S5**); cell loss being evident only at 2.5 μM NFP, akin to results with NTR 1.1^32^. This suggests that the enhanced prodrug activity of NTR 2.0 is specific to 5-nitroimidazole prodrugs (e.g. MTZ) or not relevant to 5-nitrofuran prodrugs (e.g. NFP). Of note, when higher dose assays were attempted, we observed near complete lethality with 24 h exposure at ≥5 μM NFP. This contrasts with a prior study reporting that lethality initiated at 15 μM NFP for 24 h treatments^32^.

### Bipartite transgenic NTR 2.0 expression resources

To promote dissemination of NTR 2.0 to the research community, we have created UAS and QUAS expression vectors for co-expressing NTR 2.0 with different membrane-tagged fluorescent reporters (e.g., GFP, YFP or mCherry, **Supplementary Table S8**). These plasmids also contain a “tracer” reporter (tagBFP2 for UAS lines, ECFP for QUAS lines) driven by a conserved 5’ element of *hatching enzyme* (*he*) homologs to facilitate driver-independent stock maintenance. Existing Gal4 and QF driver lines, respectively, can be combined with these resources to restrict NTR 2.0 and reporter expression to targeted cell types. Plasmids and corresponding transgenic lines will be made available through online depositories, i.e., addgene (https://www.addgene.org/) and the Zebrafish International Resource Center (ZIRC; https://zebrafish.org/home/guide.php), respectively.

## DISCUSSION

NfsB_Ec has received near-exclusive attention as a prodrug-converting NTR for both cancer gene therapy and targeted cell ablation applications^1^. However, recent evidence suggests “NTR 1.0” is a relatively inefficient NTR compared to orthologous enzymes^23,33,34^. In particular, we showed that active site residues F70 and F108 impede activity with the 5-nitroimidazole PET probe SN33623 and that rational substitution of these residues yielded generic improvements in 5-nitroimidazole reduction^23^. Despite also containing F70 and F108 residues, the naturally occurring orthologous enzyme NfsB_Vv was nearly as efficient in activating MTZ as the rationally-engineered NfsB_Ec F70A/F108Y variant **(Fig. 1e)**. Further improvement by introducing F70A/F108Y substitutions in NfsB_Vv yielded NTR 2.0, a variant with unprecedented MTZ conversion efficiency across all species tested.

First-generation (NfsB-Ec derived) NTR enzymes have enabled novel cell ablation and regeneration paradigms in the zebrafish liver^11^, pancreas^11,38^, kidney^39^, heart^11^, brain^22^, retina^40^, etc. However, the requirement for near-toxic prodrug doses to achieve effective ablation, and existence of “resistant” cell types, were major drawbacks. NTR 2.0 overcomes these limitations by enhancing MTZ-mediated ablation activity ~100-fold, eliminating the need for toxic MTZ regimens and facilitating effective ablation of resistant cells, e.g., macrophages. By facilitating cell loss at ≤1 mM MTZ, NTR 2.0 also enables prolonged cell ablation. This affords significant advantages to investigations of cell function and will allow regenerative capacities to be challenged with novel paradigms where cell ablation can be maintained for different periods of time and then released. How zebrafish respond to such challenges is unknown since mutant lines exhibiting chronic cell loss have not yet been reversed. However, results from an iterative rapid retinal cell loss paradigm suggest regenerative capacity is maintained, but with decreasing fidelity after successive bouts of cell loss and replacement^41^. The NTR 2.0/MTZ ablation system, by allowing the extent and duration of cell loss to be precisely controlled, will therefore allow regenerative capacities of discrete stem cell niches to be challenged with conditions that mimic human degenerative diseases; i.e., long-term low rates of cell loss associated with inflammatory signalling cascades.

Although NTR 2.0 exhibited improved MTZ activity across all model systems, some of the other NTR variants tested over the course of this study showed less correlation across species. In addition to the lack of activity presented here for NTR 1.1 in HEK-293 cells, we have previously encountered expression instability issues for the NfsB_Ec F70A/F108Y double mutant and other native or engineered NTR variants in HCT-116 cells^23,35–37^. Although not reported here, we have also been unable to generate stable transgenic zebrafish lines for several NTR variants that expressed well in both bacteria and HEK-293 cells. This highlights the need to test NTR variants in context when adapting to new models. The reasons for variant/context-specific expression instability of certain NTRs remain unknown, but may stem from NTR substrate promiscuity disrupting host-specific metabolic pathways. Importantly, however, we have not encountered expression stability issues for NTR 2.0 in zebrafish thus far. Nevertheless, these observations suggest heterologous expression of NTR enzymes may need to be attenuated in certain contexts.

Chronic NTR 2.0/MTZ-based cell ablation may also serve as a bona fide means of modeling degenerative diseases at the molecular level. On the face of it, the artificial nature of the system suggests there may be limited relevance to degenerative disease mechanisms. However, our recent data suggests NTR/MTZ cell ablation is mediated by DNA-damage induced cell death pathways broadly implicated in neurodegenerative disease^17^. Specifically, inhibition of Poly(ADP) ribose polymerase (PARP) protects cells from NTR/MTZ-induced ablation. PARP1 is a key element in two forms of regulated necroptotic cell death termed parthanatos and cGMP-dependent cell death. The former has been strongly linked to Parkinson’s disease^42,43^, and the latter in photoreceptor dystrophies^44^. These data are highly intriguing, emphasizing a need for further investigations of the cell death pathways elicited by the NTR/MTZ system to clarify relevance to degenerative disease mechanisms.

The possibility of applying prolonged low-dose MTZ treatments, enabled by our development of NTR 2.0, necessitates further investigations of the effects of long-term MTZ exposures. Our analysis showed no effects on survival or reproductive health after 36 days of MTZ at 0.1 or 1 mM. However, rare reports of neurotoxic effects in patients^31^ and infertility post-MTZ in rats^30^ suggest endogenous prodrug converting activities may be an issue for certain cell types. To date, studies of MTZ effects in zebrafish have been limited. One study showed 10 mM MTZ treatments altered hormone expression in the pituitary gland of larval zebrafish. Importantly, dose-response data from this group suggests treatments of ≤1 mM MTZ would not alter hormone production^29^. By enabling ablation at micromolar concentrations, NTR 2.0 therefore reduces any potential for unintended effects of MTZ.

Recently nifurpirinol (NFP) has been reported as an alternative prodrug that exhibits less variability in NTR-mediated cell ablation efficacy and is active at micromolar concentrations^32^. However, toxic effects were observed at 15 μM (24 h) and 10 μM (48 h) treatments^32^. In our hands, toxic effects arose at 5 μM NFP for 24 h. We do not know the reason for this discrepancy, but differing background strains or NFP source variation are possibilities. Nevertheless, a clear message is that because both MTZ and NFP offer only a small therapeutic index in combination with NfsB_Ec variants―i.e., are toxic at 2 to 3-fold above ablation-promoting doses―potential systemic deleterious effects remain a concern for both prodrugs when paired with first-generation NTR enzymes. While NTR 2.0 showed greatly improved activity with MTZ, it failed to show an increase in activity with NFP. This substrate specificity, however, affords an opportunity to identify NTR variants that exhibit enhanced NFP activity, but are inactive with MTZ. This may allow independent ablation of two cell types, each expressing a different NTR.

Bipartite expression systems such as Gal4/UAS^45^ and QF/QUAS^46^, as well as an ever-expanding palette of fluorescent reporters, maximize versatility of zebrafish transgenic resources and accelerate uptake of novel toolsets across the research community. Multi-reporter systems allow the dynamics of cell-cell interactions and signalling pathway activities to be visualized and quantified. Accordingly, we are creating a series of UAS and QUAS reporter/effector plasmids and transgenic lines for co-expressing various fluorescent reporters with NTR 2.0. These resources will facilitate novel NTR 2.0-enabled paradigms for interrogating cell function, exploring regenerative potential, and expanding degenerative disease modeling, thus enhancing the value of NTR targeted cell ablation system for the research community.

## Acknowledgements

This work was supported by the following grants from the National Institutes of Health, R01OD020376 (JSM and DFA) and R01NS095355 (ADL and ALC). Additional consumables support was provided by an HRC Explorer grant (contract 19/750 from the Health Research Council of New Zealand; DFA and JSM). The authors wish to thank Makeila Williams, Ben Bich, Noela Lu, and Grace Lee for technical assistance.

## Author Contributions

D.F.A. and J.S.M. conceived the research program. A.V.S., K.R.H., E.M.W., and D.F.A., designed, cloned and/or engineered all nitroreductase variants, designed bacterial experiments, p and acquired, analysed, and interpreted data. A.V.S., K.R.H., D.F.A., M.E.L-B., A.D.L., and A.L.C., designed mammalian cell culture experiments and acquired, analysed, and interpreted data. T.S.M., D.T.W., L.Z., F.M., S.N., M.T.S., and J.S.M. designed zebrafish experiments and acquired, analysed, and interpreted data. A.V.S., K.R.H., T.S.M., L.Z., S.W., K.L., D.M-L., O.L.C., generated novel transgenic zebrafish resources. D.D., and H.J. provided statistical data analysis expertise. A.V.S., T.S.M., D.F.A., and J.S.M. wrote the manuscript with input from all co-authors.

## METHODS

### Materials, bacterial strains, genes and plasmids

All reagents were purchased from Sigma-Aldrich, apart from SN33623, which was custom-synthesized by BOC Sciences. NTR candidates were PCR amplified and cloned into the *Nde*I and *Sal*I sites of two plasmids: pUCX (Addgene #60681), for biological overexpression assays, and pET28a(+) (Addgene #69864-3), for His_6_-tag protein purification. For biological assays, the *E. coli* 7NT strain was used, which bears scarless in-frame deletions of seven candidate NTR genes (*nfsA, nfsB, azoR, nemA, yieF, ycaK* and *mdaB*) and the efflux pump gene *tolC*^47^. For protein purification, the *E. coli* strain BL21(DE3) was used. The protein accession numbers for the NTRs used in this study are: NfsB_Ec = WP_000351487.1; NfsB_Vv = WP_011081872.1; NfsB_St = WP_000355874.1; NfsB_Ck = WP_047458272.1; NfsB_Kp = WP_004178896.1; NfsB_Pp = WP_010953384.1; NfsB_Cs = WP_004384956.1; FraseI_Vf = P46072.3; NfsB_Vh = WP_005435698.1; YfkO_Bs = ADE73858.1; YdgI_Bs = WP_003225379.1. Engineered gene variants were synthesized as fragments by Twist Bioscience (San Francisco, CA, USA). For expression in human cells, NTR genes were Phusion™ PCR amplified using primers that introduced a mammalian Kozak consensus sequence, a Shine-Dalgarno prokaryote consensus sequence, a TAG stop codon and Gateway™ (Invitrogen) BP recombination sites. PCR fragments were then recombined into the pDONR221™ vector (Invitrogen) using BP Clonase™ II enzyme mix (Thermo Fisher Scientific), and subsequently into the F279-V5 mammalian expression destination vector (a Gateway™ compatible plasmid which expresses inserted genes from a constitutive CMV promoter and has suppressor read through capabilities to incorporate a V5 epitope at the C-terminus)^36^ using LR Clonase™ II enzyme mix (Thermo Fisher Scientific).

For expression in zebrafish, NTR genes were PCR amplified from the pET28a+ vector using primers NfsB_Vv pTOL2_Fw and NfsB_Vv_Rv, digested with *Xma*I and *Sal*I and cloned into the *Xma*I and *Xho*I (an isoschizomer of *Sal*I) sites of a pTol2-5xUAS-GAP-tagYFP-P2A-*nfsB*_Ec vector, to swap out the parental *nfsB*_Ec for the new NTR variants. Note, the P2A element allows equimolar co-expression of the YFP reporter and NTR variants from a single transgene^48^, avoiding possible loss of NTR activity caused by fusion protein oligomerization; the GAP element is a 20 bp dual palmitoylation sequence from the zebrafish *Gap43* gene used to target reporters to the plasma membrane.

The pTol2 5xUAS:GAP-ECFP,he:GAP-ECFP plasmid was created by cutting ECFP from a 14xUAS:GAP-ECFP plasmid with *Xma*I and *BsrG*I and cloning it into the same sites to replace the 5’ EYFP reporter in the pTol2-5xUAS:GAP-EYFP,he:GAP-EYFP vector^26^. Similarly, the *hatching enzyme* promoter (*he*) driven ECFP “marker” reporter was cut from pTol2-1xUAS-GAP-tagBFP2gi,he:GAP-tagBFP2 with *Cla*I and *EcoR*V and cloned it into the same sites to replace the 3’ EYFP of the same parent vector^26^. Note, *he* promoter driven “marker” reporters provide a means of tracking germline transmission of UAS and QUAS reporter/effector transgenes in the absence of a Gal4 or QF driver, respectively. Marker reporters are strongly expressed in the hatching gland from 1-3 dpf, then fade since the hatching gland is a temporary structure^26^.

The rho:GAP-tagYFP-P2A-*nfsB*_Vv F70A/F108Y plasmid was generated by PCR amplifying the 3.7kb rhodopsin promoter from the rho:YFP-NTR plasmid using primers rho_promoter_F2 and rho_promoter_R, and cloning it into the *Fse*I and *SnaB*I sites of pTOL2-5xUAS-GAP-tagYFP-P2A-NfsB_Vv F70A-F108Y.

The pTol2 5xUAS:GAP-tagYFP-P2A-NfsB_Vv F70A/F108Y,he:GAP-tagBFP2 plasmid was generated by cutting the he:GAP-tagBFP2 element from pTol2-1xUAS-GAP-tagBFP2gi,he:GAP-tagBFP2 with *Hpa*I and *Eco*RV, and cloning it into the same sites downstream of the NTR 2.0 polyA in the vector pTol2 5xUAS:GAP-tagYFP-P2A-NfsB_Vv F70A/F108Y. The mCherry version, pTol2 5xUAS:GAP-mCherry-P2A-NfsB_Vv F70A/F108Y,he:GAP-tagBFP2, was generated by cloning the *Bsm*BI flanked portion of pTol2-5xUAS-GAP-mCherry-P2A-NTRmut,he:TagRFP into the same sites in the pTol2 5xUAS:GAP-tagYFP-P2A-NfsB_Vv F70A/F108Y,he:GAP-tagBFP2 vector. The EGFP version, pTol2 5xUAS:GAP-EGFP-P2A-NfsB_Vv F70A/F108Y,he:GAP-tagBFP2 was generated by using overlap extension PCR to fuse EGFP amplified with primers *Sbf*I-GFP_FOR and GFP_REV (from −6ascl1a:EGFP plasmid, kind gift of Dr. Dan Goldman) and 2A-NTR2.0 amplified by FW-P2A and *Kfl*I-Bglobin_REV (from the pTol2 5xUAS:GAP-tagYFP-P2A-NfsB_Vv F70A/F108Y,he:GAP-tagBFP2 vector), cutting the product with *Sbf*I and *Kfl*I and cloning it into the same sites within the pTol2 5xUAS:GAP-tagYFP-P2A-NfsB_Vv F70A/F108Y,he:GAP-tagBFP2 vector.

The pTol2 5xQUAS:GAP-tagYFP-P2A-NfsB_Vv F70A/F108Y,he:ECFP vector was created by amplifying the tagYFP to *he* promoter section of the UAS vector with the *Mlu*I-YFP_FOR and *AvrI*I-He1Pro_REV primers, digesting with *Mlu*I and *Avr*II and cloning into the same restriction sites within the pDestT2 QUAS-nls-mApple,he:ECFP plasmid (kindly provided by Marnie Halpern). All relevant primers are listed in **Supplementary Table S9**. Full sequences of NTR variants, vectors, and transgenes used to create new lines are provided in **Supplementary Data File S10**.

### Creation of NTR-expressing and fluorescent HEK-293 cell lines

Plasmid encoding a nitroreductase (F279-V5:*ntr*, where *ntr* represents a gene encoding NTR 1.0, NTR 1.1, NfsB_Vv, or NTR 2.0), or a fluorescent protein (pCDNA3-GFP; Addgene plasmid #74165 or mCherry2-N1; Addgene plasmid #54517) was used to transfect HEK-293 cells at 70-90% confluency using Lipofectamine 3000 reagent (Thermo Fisher Scientific) as per manufacturer’s instructions. To generate stable polyclonal cell lines, 48 h following transfection the medium in transfected wells was exchanged for medium supplemented with the appropriate selection antibiotic. Cells which had stably integrated plasmid DNA were selected for by multiple passage cycles in medium containing escalating concentrations of the selection antibiotic (1-3 μM in the case of puromycin, or 500-900 μg/ml in the case of G418) until cell death was no longer evident. Selection antibiotic concentrations were determined following generation of a dose response curve with wild-type cells.

### Bacterial cytotoxicity assays

Cells of *E. coli* 7NT pUCX:*nitroreductase* strains were used to inoculate wells of a sterile 96 well microplate containing 200 μl LB supplemented with antibiotic and 0.2% (w/v) glucose. Cultures were incubated overnight at 30 °C, 200 rpm and the following morning, cultures were diluted 20-fold into 15 ml centrifuge tubes containing 2 ml induction medium (LB supplemented with antibiotic, 0.2% (w/v) glucose and 50 mM IPTG). Cultures were incubated at 30 °C, 200 rpm for 2.5 h. Aliquots (30 μl) of each culture were added to wells of a sterile 384 well microplate containing 30 μl induction medium ± two-fold the final prodrug concentration. Each culture was exposed, in duplicate, to 15 conditions containing 2- or 1.5-fold increasing titrations of prodrug and one induction medium-only control. The medium was supplemented with DMSO as appropriate, ensuring that the final DMSO concentration remained <4%. OD_600_ readings were recorded using an EnSpire™ 2300 Multilabel Reader (PerkinElmer). It was ensured that the starting OD_600_ values across strains were comparable and fell within the range of 0.12-0.18. Cultures were incubated at 30 °C, 200 rpm, for a further 4 h, after which OD_600_ readings were recorded once more. The increase in OD_600_ of corresponding challenged and unchallenged wells for each strain were compared and used to calculate percentage growth at each prodrug concentration. EC_50_ values (the concentration of prodrug at which growth was reduced by 50% relative to the unchallenged control) were calculated using a dose-response inhibition four-parameter variable slope equation in GraphPad Prism 7.0 (GraphPad Software Inc).

### Mammalian cytotoxicity assays

A standard 48 h MTS (3-(4,5-dimethylthiazol-2-yl)-5-(3-carboxymethoxyphenyl)-2-(4-sulfophenyl)-2H-tetrazolium) cell viability assay was used to evaluate NTR-mediated activation of MTZ and associated cytotoxic activity against the human embryonic kidney cell line HEK-293 (n ≥ 3 independent experiments with duplicate wells per experiment). Cell lines stably expressing an NTR were treated with MTZ at various concentrations, and a dose–response was generated relative to a control of untreated HEK-293 cells. HEK-293 cell lines were seeded in 100 μL aliquots into 6.25 mm diameter culture wells at a density of 180,000 cells/mL. Cells were seeded in RPMI medium supplemented with 1× glutaMAX, 10% fetal calf serum and 1% penicillin/streptomycin. Cells were left to adhere in a humidified incubator at 37°C, 5% CO_2_ for approximately 16 hours before being challenged with 100 μL of RPMI medium supplemented with 1× glutaMAX ± 2× final MTZ concentration. Cells were incubated at 37°C, 5% CO_2_ for 48 h. Following challenge, 10 μL of CellTitre 96 Aqueous One Solution Cell Proliferation Assay reagent (Promega; Madison, USA) was added to each well (as well as to a medium-only containing well to allow for baseline medium absorbance subtraction) and cells were incubated for a further hour at 37 °C, 5% CO_2_. Absorbance of wells was measured at 490 nm, and the absorbance value of the media-only control well was subtracted from all other measurements. The baseline-subtracted absorbance of challenged wells was compared to that of an unchallenged control well to determine percentage cell viability for each MTZ concentration. EC_50_ values were calculated using a dose-response inhibition four-parameter variable slope equation in GraphPad Prism 7.0 (GraphPad Software Inc; La Jolla, USA). Calculated EC_50_ values are the averages of at least three biological replicates.

CHO-KI cells were grown in Ham’s F12 medium supplemented with 10% FBS and pen/strep. To obtain cultures for prodrug killing experiments, cells were plated on 10 cm plates and, when 50-60% confluent, transfected with either the NTR 1.1-mCherry construct (mCherry-P2A-NfsB_Ec T41Q/N71S/F124T) or the NTR 2.0-YFP construct (tagYFP-P2A-NfsBVv F70A/F108Y), cloned into pcDNA3). Transfection was performed using Lipofectamine 3000 according to manufacturer’s instructions. Stable clones were generated by adding G418 (0.5 mg/ml) to growth medium and selecting resistant colonies that exhibited mCherry or YFP fluorescence. Colonies were picked manually, but since not all cells within a colony expressed a given reporter (mCherry or YFP), colonies were re-cloned by limiting dilution. Cells were maintained in G418 selection medium until plated for MTZ treatment and MTT (3-[4, 5-dimethylthiazol-2-yl]-2, 5 diphenyl tetrazolium bromide) cell viability assays.

For MTZ dose-response assays, transfected CHO-KI cells were plated into 96-well plates at a concentration of 10,000 cells per well in 100 μl of medium, without G418, and allowed to adhere overnight. On the following day, medium was carefully removed and replaced with 100 μl of medium containing varying concentrations of MTZ (a minimum of 4 wells per MTZ concentration were tested). CHO-KI cells expressing NTR1.1-mCherry were treated with MTZ (0-10 mM) solubilized in water; control wells (0 MTZ) contained 10% water in growth medium. CHO-KI cells expressing NTR 2.0-YFP were treated with MTZ (0-1 mM) solubilized in 0.1% DMSO; control wells (0 MTZ) contained 0.1% DMSO in growth medium. After 24 h of growth in MTZ, the medium in each well was replenished by adding an additional 100 μl medium containing the appropriate concentration of MTZ. Following 48 h total of MTZ treatment, MTT assays (adapted from Mossman, 1983^25^) were performed to assess cell viability: 20 μL of 5 mg/ml of MTT (Thiazolyl Blue Tetrazolium Bromide, Sigma-Aldrich, M5655) in Dulbecco’s PBS was added to each well. The plates were incubated for 3-4 h in a humidified 37°C CO_2_ incubator. Following incubation, the supernatant was removed, and 100 μL of DMSO was added to each culture well and mixed. The plates were incubated for 30 min more at 37°C/5% CO_2_, following which absorbance was read at 560 nm on a plate reader. The absorbance values of wells with DMSO alone were used to establish the average background absorbance value, which was subtracted from the average absorbance obtained for each MTZ treatment condition. Cell viability (%) was calculated by dividing the average absorbance for each MTZ treatment condition by the absorbance value for cells treated with vehicle alone, multiplied by 100. EC_50_ values were calculated using GraphPad Prism 8.0 software (Graphpad Software Inc) using the log(inhibitor) vs response (variable slope--4 parameters) function.

### Nitroreductase expression and purification

A 3 ml culture of LB medium supplemented with kanamycin was inoculated with BL21 (DE3) cells expressing a pET28a+:*nitroreductase* construct and grown overnight at 37 °C, 200 rpm. The overnight culture was diluted 100-fold into 100 ml LB medium containing 50 μg.ml^−1^ kanamycin and incubated at 37 °C, 200 rpm until an OD_600_ of 0.6-0.8 was reached. The culture was chilled on ice for 20 min after which IPTG was added to a final concentration of 0.5 mM to induce nitroreductase expression. The culture was incubated for a further 24 h at 18 °C and the cells were pelleted by centrifugation (2600 x g, 4 °C, 20 min). The cell pellet was lysed by resuspension in Bugbuster® (Novagen) (4 ml for every 50 ml of pre-pelleted culture) and incubated at room temperature on an orbital shaker for 20 min. The cell lysate was centrifuged at 17,000 x g, 4 °C for 15 min to achieve separation of the insoluble and soluble fractions. Soluble fractions were decanted and placed on ice. Purification of His_6_-tagged proteins from the soluble fraction was achieved using Ni-NTA His.Bind® Resin (Merck), a peristaltic pump and a series of column washes with buffers of increasing imidazole content. Ni-NTA His.Bind® Resin (2-4 ml) was packed into a 10 ml Pierce disposable column (Thermo Fisher) and connected to the peristaltic pump. The resin was washed with 4 ml of ddH_2_O and charged by flowing through 8 ml of Charge buffer (50 mM NiSO_4_). The soluble fraction of the lysed cells was pumped through the column, followed by 10 ml Bind buffer (500 mM NaCl, 20 mM Tris-HCl pH 7.9, 5 mM imidazole) and 8 ml Wash buffer (500 mM NaCl, 20 mM Tris-HCl pH 7.9, 60 mM imidazole). The pump tubing was evacuated, after which 4 ml of Elution buffer (500 mM NaCl, 20 mM Tris-HCl pH 7.9, 1 M imidazole) was run through the column and flow through was collected in 500 μl fractions. The three most yellow fractions (indicative of the highest concentration of FMN-bound nitroreductase) were pooled. Resin beads were washed and stored in Strip buffer (500 mM NaCl, 20 mM Tris-HCl pH 7.9, 100 mM EDTA) at 4 °C. Pooled nickel-purified protein fractions were incubated with excess FMN cofactor (1 mM) for 1 h on ice. Buffer-exchange into 40 mM Tris-Cl pH 7.0 was carried out using a 5 ml HiTrap™ desalting column (GE Healthcare) as per the manufacturer’s instructions. Collected flow-through was restricted to the first 1.5 ml, as flow-through beyond 1.5 ml was likely to contain contaminating free FMN.

### Steady-state enzyme kinetics

Michaelis-Menten kinetic parameters were derived using NADPH as a cofactor in the presence of a NADPH regenerating system (*Bacillus subtilis* glucose dehydrogenase + glucose) to maintain constant NADPH levels. Where possible, substrate concentrations were varied from approximately 0.2× *K*_*M*_ to 5× *K*_*M*_ however spectroscopic absorption limits prevented concentrations above 3 mM MTZ from being accurately measured. Reactions were prepared in a 1 mm pathlength quartz cuvette in a total volume of 100 μl containing 100 μM NADPH, 5 mM glucose, 0.55 μM *B. subtilis* glucose dehydrogenase, 10 μM NTR, 50 mM sodium phosphate buffer pH 7.0 and a varying concentration of MTZ. The absorbance of each reaction at 340 nm was followed, and mean rates of reduction at each MTZ concentration were obtained from a minimum of triplicate repeats. The extinction coefficient of MTZ reduction at 340 nm was calculated at 1,980 M^−1^cm^−1^ by measuring the drop in absorbance of 100 μM MTZ following complete reduction by NfsA_Ec (after the innate oxidase activity of this enzyme had consumed residual NADPH). Non-linear regression analysis and Michaelis-Menten curve fitting were performed using Graphpad Prism 8.0 (Graphpad Software Inc). Raw Michaelis-Menten plots are presented as **Supplementary Fig. S2**.

### Fluorescence live cell imaging

For cultured cells, brightfield and fluorescence images were taken on an Olympus IX51 inverted microscope and merged to allow visualization of the ratio of fluorescent to non-fluorescent cells. Confocal intravital imaging of zebrafish was performed using an upright Olympus FV1000 confocal microscope and dipping cone objectives (20× water immersion, NA 0.95 and 4× air, NA 0.3).

### Zebrafish husbandry and transgenic resources

Zebrafish were maintained under standard growth conditions at 28.5 °C in a 10 h dark / 14 h light cycle. All zebrafish were treated in accordance with approved protocols reviewed by the Johns Hopkins Animal Care and Use Committee. Unless otherwise noted all zebrafish used were in the *roy orbison* (*roy; mpv17*^*a9/a9*^) mutant background and treated with 200 nM phenylthiourea (PTU) at 16-24 hours post-fertilization (hpf) to facilitate live imaging and quantification of fluorescent reporters. New zebrafish transgenic lines were created by co-injecting miniTol2-based expression plasmids (cloning details above) and 25 pg of Tol2 mRNA into one-cell stage zebrafish embryos and screening for germline transmission. UAS reporter lines, *Tg(5xUAS:GAP-tagYFP-P2A-nfsB_Vv F70A/F108Y)jh513* and *Tg(5xUAS:GAP-ECFP,he:GAP-ECFP)gmc1913*, were created by injecting into eggs from the *Et(2xNRSE-fos:KALTA4)gmc617* driver line to allow visualization of reporter expression. The *Tg(rho:GAP-tagYFP-P2A-nfsB_Vv F70A/F108Y)jh405* line was created by injecting into *roy*^*a9/a9*^ eggs. A full list of all zebrafish transgenic lines used or created here, as well as abbreviations, delineation of NTR variant, and references is provided in **Supplementary Table S10**.

### Automated reporter quantification *in vivo* (ARQiv)

A plate reader-based method for *in vivo* reporter quantification was performed as previously detailed^15,16^. Briefly, zebrafish were aliquoted individually into wells of 96-well, black, polypropylene, U-bottom plates containing 300 μL of embryo media (+200 nM PTU) and the indicated concentration of MTZ or NFP. On the day of analysis, larvae were anesthetized by addition of 50 μL clove oil (350 ppm) for 15 minutes. A Tecan Infinite M1000 plate reader was then used to quantify fluorescence levels in individual fish. Nine regions per well were scanned to account for random orientation of zebrafish using excitation/emission and bandwidth settings of 508 ±5 nm and 528 ±5 nm, respectively, for tagYFP; and 514 ±5 nm and 538 ±10 nm, respectively, for EYFP. The data were processed by first defining a fluorescence “signal” cutoff as the averaged maximal values, plus three standard deviations, of non-transgenic controls. All regional scan values greater than or equal to the signal cutoff were summed to calculate the fluorescence level of each sample. The data were then normalized to a “signal window” bounded by the non-transgenic controls (set at 0%) and 0 MTZ controls (set at 100%) by subtracting the averaged maximal value of the non-transgenic controls from all summed signal values and then dividing the resulting values by the averaged signal from the 0 MTZ controls. Absolute effect sizes for all experimental conditions are expressed as a percentage of the normalized signal window for each assay to facilitate comparisons across biological repeats.

### Statistical Analysis of ARQiv and IMARIS data

A custom R code (R v3.6.3 and R Studio v1.2.5033) was written for statistical analysis of the data. To prepare the data for analysis, normalized signal values were organized into columns per each treatment condition in Excel. The R code then subjected treatment groups to pairwise comparisons to either the non-ablated (0 mM prodrug) or non-transgenic controls using Welch’s unequal variances t-test (t.test() function). The R code produced the following data points: absolute effect sizes (mean differences between the treatment group and controls), 95% confidence intervals, degrees of freedom, sample sizes, and *p*-values. All p-values were then adjusted for multiple comparisons using the p.adjust() function, resulting in Bonferroni-corrected *p’*-values.

### Microglia/macrophage ablation

Transgenic *mpeg:NTR1.1-YFP* or double transgenic *mpeg:Gal4;UAS:YFP-NTR2.0* larvae were treated with 01, 1, or 10 mM MTZ for 48 h from 5-7 dpf. Intravital imaging performed on pre- and post-treatment larvae were acquired with an Olympus FV1000. Quantification of immune cell number was performed in IMARIS (v9.2; Bitplane) from z-stacks rendered with FiJi (i.e., ImageJ v1.52h; NIH). Statistical analyses were performed as described above.

### Long-term MTZ exposure

Late larval non-transgenic *roy orbison* (*roy*^*a9/a9*^) fish (15 dpf) were exposed to 0, 0.1, 1 or 10 mM MTZ for 36 days (until 51 dpf). MTZ treatments were initiated at 15 dpf to avoid increases in larval death typical of late larval *roy* mutant development, i.e., between 10 and 12 dpf. Groups of 30 to 33 larvae were maintained in 1 L of stagnant system water at the indicated concentrations of MTZ, with 750 mL of water ±MTZ replaced every 2-3 days, and the experiment repeated three times. To assess fecundity post-MTZ treatment, 12-15 pairwise in-crosses of 3 month old fish from each treatment group were setup on three separate occasions. From each successful mating, the number of embryos laid, fertilization rates and offspring survival were recorded until 5 dpf.

### Statistical Analysis of Long-term MTZ exposure data

Juvenile survival data was subjected to Log-rank (Mantel-Cox) tests and Gehan-Breslow-Wilcoxon tests (Prism, GraphPad) to generate chi-square and *p*-values. All other data was subjected to ANOVA followed by Dunnett’s multiple comparisons test, depending on the outcome of Levene’s test, to generate 95% confidence intervals and *p*-values. All *p*-values were subsequently adjusted for multiple comparisons with Bonferroni correction. Absolute effect sizes were calculated as the mean difference between control and experimental conditions.

## REFERENCES

1. Williams, E. M. et al. Nitroreductase gene-directed enzyme prodrug therapy: insights and advances toward clinical utility. Biochem. J. 471, 131–153 (2015).

2. Roldán, M. D., Pérez-Reinado, E., Castillo, F. & Moreno-Vivián, C. Reduction of polynitroaromatic compounds: the bacterial nitroreductases. FEMS Microbiol. Rev. 32, 474–500 (2008).

3. Venitt, S. & Crofton-Sleigh, C. The toxicity and mutagenicity of the anti-tumour drug 5-aziridino-2,4-dinitrobenzamide (CB1954) is greatly reduced in a nitroreductase-deficient strain of E. coli. Mutagenesis 2, 375–81 (1987).

4. Knox, R. J. et al. The nitroreductase enzyme in Walker cells that activates 5-(aziridin-1-yl)-2,4-dinitrobenzamide (CB 1954) to 5-(aziridin-1-yl)-4-hydroxylamino-2-nitrobenzamide is a form of NAD(P)H dehydrogenase (quinone) (EC 1.6.99.2). Biochem. Pharmacol. 37, 4671–7 (1988).

5. Lewis, K. Platforms for antibiotic discovery. Nat. Rev. Drug Discov. 12, 371–387 (2013).

6. Bridgewater, J. A. et al. Expression of the bacterial nitroreductase enzyme in mammalian cells renders them selectively sensitive to killing by the prodrug CB1954. Eur. J. Cancer 31A, 2362–2370 (1995).

7. Drabek, D., Guy, J., Craig, R. & Grosveld, F. The expression of bacterial nitroreductase in transgenic mice results in specific cell killing by the prodrug CB1954. Gene Ther. 4, 93–100 (1997).

8. Clark, A. J. et al. Selective cell ablation in transgenic mice expression E. coli nitroreductase. Gene Ther. 4, 101–10 (1997).

9. Bridgewater, J. A., Knox, R. J., Pitts, J. D., Collins, M. K. & Springer, C. J. The bystander effect of the nitroreductase/CB1954 enzyme/prodrug system is due to a cell-permeable metabolite. Hum. Gene Ther. 8, 709–717 (1997).

10. Medico, E., Gambarotta, G., Gentile, A., Comoglio, P. M. & Soriano, P. A gene trap vector system for identifying transcriptionally responsive genes. Nat. Biotechnol. 19, 579–82 (2001).

11. Curado, S. et al. Conditional targeted cell ablation in zebrafish: a new tool for regeneration studies. Dev. Dyn. 236, 1025–35 (2007).

12. White, D. T. & Mumm, J. S. The nitroreductase system of inducible targeted ablation facilitates cell-specific regenerative studies in zebrafish. Methods 62, 232–240 (2013).

13. Ariga, J., Walker, S. L. & Mumm, J. S. Multicolor time-lapse imaging of transgenic zebrafish: visualizing retinal stem cells activated by targeted neuronal cell ablation. J. Vis. Exp. 43, e2093 (2010).

14. White, D. T. et al. Immunomodulation-accelerated neuronal regeneration following selective rod photoreceptor cell ablation in the zebrafish retina. Proc. Natl. Acad. Sci. 114, E3719–E3728 (2017).

15. Walker, S. L. et al. Automated reporter quantification in vivo: High-throughput screening method for reporter-based assays in zebrafish. PLoS One 7, e29916 (2012).

16. White, D. T. et al. ARQiv-HTS, a versatile whole-organism screening platform enabling in vivo drug discovery at high-throughput rates. Nat. Protoc. 11, 2432–2453 (2016).

17. Zhang, L. et al. Large-scale Phenotypic Drug Screen Identifies Neuroprotectants in Zebrafish and Mouse Models of Retinitis Pigmentosa. bioRxiv 2020.03.26.010009 (2020). doi:10.1101/2020.03.26.010009

18. Mathias, J. R., Zhang, Z., Saxena, M. T. & Mumm, J. S. Enhanced cell-specific ablation in zebrafish using a triple mutant of Escherichia coli nitroreductase. Zebrafish 11, 85–97 (2014).

19. Godoy, R., Noble, S., Yoon, K., Anisman, H. & Ekker, M. Chemogenetic ablation of dopaminergic neurons leads to transient locomotor impairments in zebrafish larvae. J. Neurochem. 135, 249–260 (2015).

20. Petrie, T. A. et al. Macrophages modulate adult zebrafish tail fin regeneration. Development 141, 2581–91 (2014).

21. Guise, C. P., Grove, J. I., Hyde, E. I. & Searle, P. F. Direct positive selection for improved nitroreductase variants using SOS triggering of bacteriophage lambda lytic cycle. Gene Ther. 14, 690–8 (2007).

22. Tabor, K. M. et al. Direct activation of the Mauthner cell by electric field pulses drives ultrarapid escape responses. J. Neurophysiol. 112, 834–44 (2014).

23. Williams, E. M. et al. Engineering *Escherichia coli* NfsB To Activate a Hypoxia-Resistant Analogue of the PET Probe EF5 To Enable Non-Invasive Imaging during Enzyme Prodrug Therapy. Biochemistry 58, 3700–3710 (2019).

24. Cory, A. H., Owen, T. C., Barltrop, J. A. & Cory, J. G. Use of an Aqueous Soluble Tetrazolium/Formazan Assay for Cell Growth Assays in Culture. Cancer Commun. 3, 207–212 (1991).

25. Mosmann, T. Rapid colorimetric assay for cellular growth and survival: Application to proliferation and cytotoxicity assays. J. Immunol. Methods 65, 55–63 (1983).

26. Xie, X. et al. Silencer-delimited transgenesis: NRSE/RE1 sequences promote neural-specific transgene expression in a NRSF/REST-dependent manner. BMC Biol. 10, 93 (2012).

27. Distel, M., Wullimann, M. F. & Köster, R. W. Optimized Gal4 genetics for permanent gene expression mapping in zebrafish. Proc. Natl. Acad. Sci. U. S. A. 106, 13365–70 (2009).

28. Davison, J. M. et al. Transactivation from Gal4-VP16 transgenic insertions for tissue-specific cell labeling and ablation in zebrafish. Dev. Biol. 304, 811–24 (2007).

29. Cheng, X. et al. Effects of metronidazole on proopiomelanocortin a gene expression in zebrafish. Gen. Comp. Endocrinol. 214, 87–94 (2015).

30. McClain, R. Effect of metronidazole on fertility and testicular function in male rats. Fundam. Appl. Toxicol. 12, 386–396 (1989).

31. Sørensen, C. G., Karlsson, W. K., Amin, F. M. & Lindelof, M. Metronidazole-induced encephalopathy: a systematic review. Journal of Neurology 267, 1–13 (2020).

32. Bergemann, D. et al. Nifurpirinol: A more potent and reliable substrate compared to metronidazole for nitroreductase-mediated cell ablations. Wound Repair Regen. 26, 238–244 (2018).

33. Williams, E. M. et al. A cofactor consumption screen identifies promising NfsB family nitroreductases for dinitrotoluene remediation. Biotechnol. Lett. 41, 1155–1162 (2019).

34. Crofts, T. S. et al. Discovery and Characterization of a Nitroreductase Capable of Conferring Bacterial Resistance to Chloramphenicol. Cell Chem. Biol. 26, 559–570.e6 (2019).

35. Copp, J. N. et al. Engineering a Multifunctional Nitroreductase for Improved Activation of Prodrugs and PET Probes for Cancer Gene Therapy. Cell Chem. Biol. 24, 391–403 (2017).

36. Prosser, G. A. et al. Creation and screening of a multi-family bacterial oxidoreductase library to discover novel nitroreductases that efficiently activate the bioreductive prodrugs CB1954 and PR-104A. Biochem. Pharmacol. 85, 1091–103 (2013).

37. Swe, P. M. et al. Targeted mutagenesis of the Vibrio fischeri flavin reductase FRase I to improve activation of the anticancer prodrug CB1954. Biochem. Pharmacol. 84, 775–83 (2012).

38. Pisharath, H., Rhee, J. M., Swanson, M. a, Leach, S. D. & Parsons, M. J. Targeted ablation of beta cells in the embryonic zebrafish pancreas using E. coli nitroreductase. Mech. Dev. 124, 218–29 (2007).

39. Huang, J. et al. A zebrafish model of conditional targeted podocyte ablation and regeneration. Kidney Int. 83, 1193–200 (2013).

40. Montgomery, J. E., Parsons, M. J. & Hyde, D. R. A novel model of retinal ablation demonstrates that the extent of rod cell death regulates the origin of the regenerated zebrafish rod photoreceptors. J. Comp. Neurol. 518, 800–814 (2010).

41. Ranski, A. H., Kramer, A. C., Morgan, G. W., Perez, J. L. & Thummel, R. Characterization of retinal regeneration in adult zebrafish following multiple rounds of phototoxic lesion. PeerJ 6, e5646 (2018).

42. Lee, Y. et al. Parthanatos mediates AIMP2-activated age-dependent dopaminergic neuronal loss. Nat. Neurosci. 16, 1392–1400 (2013).

43. Kam, T.-I. et al. Poly(ADP-ribose) drives pathologic α-synuclein neurodegeneration in Parkinson’s disease. Science 362, eaat8407 (2018).

44. Power, M. et al. Cellular mechanisms of hereditary photoreceptor degeneration - Focus on cGMP. Prog. Retin. Eye Res. 74, 100772 (2020).

45. Halpern, M. E. et al. Gal4/UAS transgenic tools and their application to zebrafish. Zebrafish 5, 97–110 (2008).

46. Subedi, A. et al. Adoption of the Q transcriptional regulatory system for zebrafish transgenesis. Methods 66, 433–40 (2014).

47. Copp, J. N., Hanson-Manful, P., Ackerley, D. F. & Patrick, W. M. Error-prone PCR and effective generation of gene variant libraries for directed evolution. Methods Mol. Biol. 1179, 3–22 (2014).

48. Provost, E., Rhee, J. & Leach, S. D. Viral 2A peptides allow expression of multiple proteins from a single ORF in transgenic zebrafish embryos. Genesis 45, 625–629 (2007).

49. Ellett, F., Pase, L., Hayman, J. W., Andrianopoulos, A. & Lieschke, G. J. Mpeg1 Promoter Transgenes Direct Macrophage-Lineage Expression in Zebrafish. Blood 117, e49–56 (2011).

