## Supplemental Materials for "NTR 2.0: a rationally-engineered prodrug converting enzyme with substantially enhanced efficacy for targeted cell ablation"

### SUPPLEMENTARY MATERIALS

### Supplementary Fig. S1: Identification of ‘SN33623-blocking’ residues in NfsB_Ec-like nitroreductases.

NfsB_Ec 68 YV**F**NERKMLDASHVVVFCAKTAMDDVWLKLVVDQEDADGR**F**AT 110

NfsB_Vv 68 FA**F**NERKILDASHVVVFCAKTSIDEAYLLDLLQSEDKDGR**F**AD 110

NfsB_Kp 68 YV**F**NERKILDASHVVVFCAKTAMDDAWLQRVVDQEEADGR**F**AT 110

NfsB_Ck 68 FV**F**NERKMLDASHVVVFCAKTAMDDAWLDRVVDQEDADGR**F**AT 110

NfsB_St 68 YT**F**NERKMLDASHVVVFCAKTAMDDAWLQRVVDQEDADGR**F**AT 110

NfsB_Pp 68 YA**Y**NTQKLLDASHVIVFCTRTEMTEEHLNAVLDQEAADGR**F**RD 110

```
:. * :*:******:***::* : :  *  :::.*  ***:
```

Partial protein alignment of NfsB_Ec and NfsB_Ec-like enzymes (residues 68 – 110) with ‘SN33623-blocking’ residues highlighted in red. Identical (*), conservative (:), and semi-conservative (.) amino acid differences are indicated Alignment performed using Clustal Omega sequence alignment tool (default settings; *http://www.ebi.ac.uk/Tools/msa/clustalo*).

### Supplementary Fig. S2: Michaelis-Menten reaction curves of purified nitroreductase variants with metronidazole.

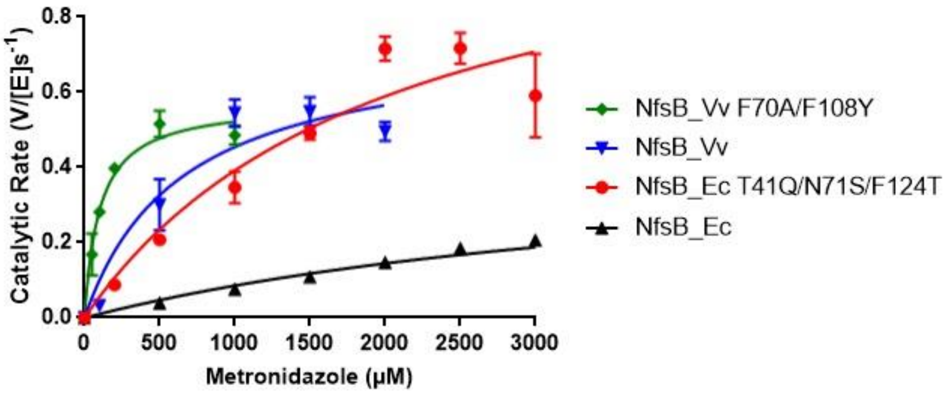


The indicated NTR enzymes were purified and assayed for MTZ conversion activity across a concentration range spanning from *ca*. 0.2× to 5× the $K_M$ for each variant and at a minimum of three repeats per condition. Error bars indicate ± SD.

| Nitroreductase | $k_{cat}$ ($s^{-1}$)* | | | $K_M$ (µM)* | | | $k_{cat}/K_M$ ($mM^{-1}$ $s^{-1}$) | | |
|---|---|---|---|---|---|---|---|---|---|
| NfsB_Ec | 0.47 | ± | 0.12 | 4400 | ± | 1800 | 110 | ± | 51 |
| NfsB_Ec T41Q/N71S/F124T | 1.19 | ± | 0.35 | 2000 | ± | 1200 | 580 | ± | 380 |
| NfsB_Vv | 0.75 | ± | 0.17 | 650 | ± | 410 | 1200 | ± | 770 |
| NfsB_Vv F70A/F108Y | 0.57 | ± | 0.03 | 100 | ± | 21 | 5700 | ± | 1200 |

**Supplementary Table S1: Michaelis-Menten kinetic parameters for the reduction of metronidazole by purified His$_6$-tagged nitroreductases.** Reduction rates at varying MTZ concentrations were measured at a fixed steady-state concentration of NADPH (100 µM) in the presence of a cofactor regeneration system. MTZ reduction was monitored at 340 nm. At each substrate concentration, data was obtained from a minimum of triplicate repeats, and error indicates ± SD. $K_M$ and $k_{cat}$ values were derived using GraphPad Prism 7*.0 (GraphPad Software Inc)*. The asterisk (*) indicates that these are apparent kinetic parameters as measured at 100 µM NADPH. An extinction coefficient of 1,980 $M^{-1}cm^{-1}$ was used to account for the loss of absorbance at 340 nm due to MTZ reduction.

### Supplementary Fig. S3: Dose-response assays of nitroreductase variants with metronidazole in vertebrate cells.

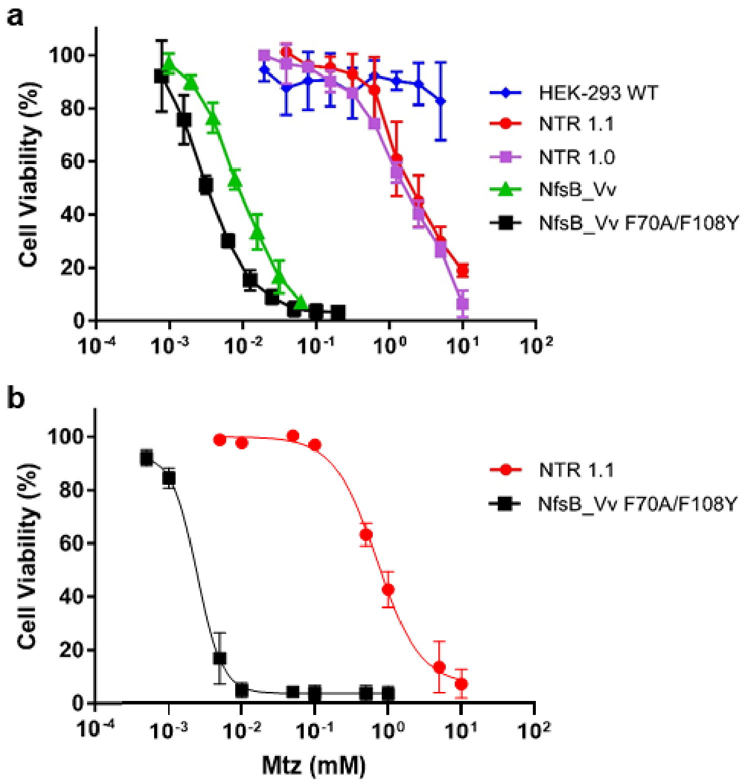


**a,b,** NTR variants were stably expressed in HEK-293 **(a)** or CHO-K1 cells **(b)** and tested across the indicated MTZ concentrations. Survival rates were obtained by MTS **(a)** or MTT **(b)** assays, from a minimum of triplicate biological repeats, error bars indicate ± 1 SD.

| **HEK-293 cell line** (MTS assay) | **Metronidazole $EC_{50}$ (µM)** |
|---|---|
| **NTR 1.1.** (NfsB_Ec T41Q/N71S/F124T) | 2300 ± 970 |
| **NTR 1.0** (NfsB_Ec) | 1700 ± 200 |
| **NfsB_Vv** | 9 ± 2 |
| **NfsB_Vv F70A/F108Y** | 3 ± 1 |
| **HEK-293 WT** | >5000 |
| **CHO-K1 cell line** (MTT assay) | **Metronidazole $EC_{50}$ (µM)** |
| **NTR 1.1** (NfsB_Ec T41Q/N71S/F124T) | 690 ± 66 |
| **NfsB_Vv F70A/F108Y** | 4 ± 2 |

**Supplementary Table S2: Metronidazole $EC_{50}$ values for mammalian stable cell lines over-expressing candidate nitroreductase variants.** HEK-293 and CHO-K1 cells over-expressing the indicated NTR enzymes were challenged with a 2-fold dilution series of MTZ (1 – 1000 µM) and incubated for 48 h prior to the addition of MTS or MTT reagent, respectively. The absorbance of MTZ-challenged wells ($OD_{490}$ nm) was compared to unchallenged control wells. Percent viability relative to unchallenged control was calculated for each drug concentration and GraphPad Prism 7.0 used to calculate $EC_{50}$ values. Data presented are the average of at least three biological replicates ± SD.

Supplementary Fig. S4: Neuronally-restricted co-expression of YFP and NTR 2.0.

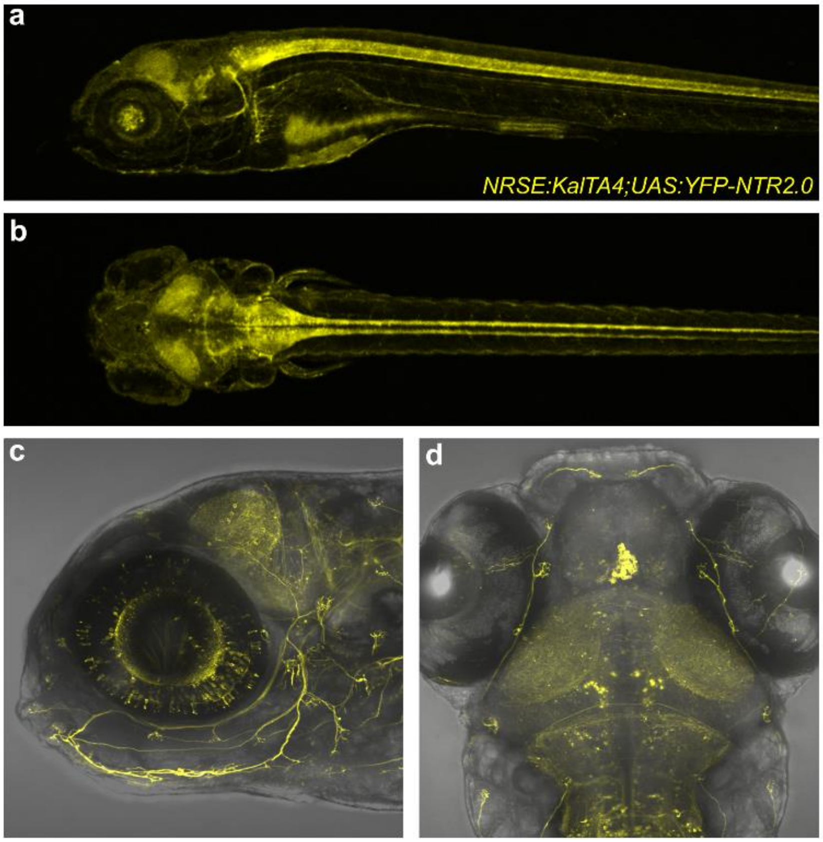


**a-d,** Intravital confocal images of *NRSE:KalTA4;UAS:YFP-NTR2.0* double transgenic fish illustrating neuronally-restricted expression of a membrane-tagged YFP reporter co-expressed with NTR 2.0. Images were taken at 5 dpf from lateral **(a,c)** and dorsal perspectives **(b, d)**.

### Statistical analysis of 24 h MTZ titration (Fig. 3a)

| Compared to 0 mM MTZ | | 12.5 µM | 25 µM | 50 µM | 100 µM | 200 µM | Non-Tg |
|---|---|---|---|---|---|---|---|
| **effect size** | | -0.15 | -0.23 | -0.70 | -0.87 | -0.96 | -1.00 |
| **95%CI lower** | | -0.33 | -0.42 | -0.84 | -1.00 | -1.08 | -1.12 |
| **95%CI upper** | | 0.02 | -0.03 | -0.55 | -0.74 | -0.84 | -0.88 |
| ***p′*-value** | | 0.532 | 0.143 | $3.4^{E-16}$ | $1.2^{E-26}$ | $6.8^{E-30}$ | $2.8^{E-31}$ |
| **N** | 116 | 50 | 80 | 78 | 122 | 50 | 140 |

| Compared to Non-Tg | | 0 µM | 12.5 µM | 25 µM | 50 µM | 100 µM | 200 µM |
|---|---|---|---|---|---|---|---|
| **effect size** | | 1.00 | 0.85 | 0.77 | 0.30 | 0.13 | 0.04 |
| **95%CI lower** | | 0.88 | 0.72 | 0.62 | 0.22 | 0.09 | 0.03 |
| **95%CI upper** | | 1.12 | 0.98 | 0.93 | 0.39 | 0.17 | 0.05 |
| ***p′*-value** | | $2.8^{E-31}$ | $9.2^{E-17}$ | $1.7^{E-14}$ | $1.1^{E-08}$ | $1.0^{E-08}$ | $8.6^{E-11}$ |
| **N** | 140 | 116 | 50 | 80 | 78 | 122 | 50 |

### Statistical analysis of 48 h MTZ titration (Fig. 3b)

| Compared to 0 mM MTZ | | 12.5 µM | 25 µM | 50 µM | 100 µM | 200 µM | Non-Tg |
|---|---|---|---|---|---|---|---|
| **effect size** | | -0.36 | -0.71 | -0.83 | -0.91 | -0.97 | -1.00 |
| **95%CI lower** | | -0.56 | -0.87 | -0.97 | -1.06 | -1.11 | -1.14 |
| **95%CI upper** | | -0.15 | -0.55 | -0.68 | -0.77 | -0.83 | -0.86 |
| ***p′*-value** | | 0.005 | $1.2^{E-14}$ | $2.6^{E-20}$ | $1.6^{E-23}$ | $2.1^{E-25}$ | $3.0^{E-26}$ |
| **N** | 110 | 53 | 99 | 106 | 104 | 59 | 121 |

| Compared to Non-Tg | | 0 µM | 12.5 µM | 25 µM | 50 µM | 100 µM | 200 µM |
|---|---|---|---|---|---|---|---|
| **effect size** | | 1.00 | 0.64 | 0.29 | 0.17 | 0.09 | 0.03 |
| **95%CI lower** | | 0.86 | 0.49 | 0.21 | 0.12 | 0.05 | 0.01 |
| **95%CI upper** | | 1.14 | 0.80 | 0.38 | 0.22 | 0.12 | 0.05 |
| ***p′*-value** | | $3.0^{E-26}$ | $3.4^{E-10}$ | $4.3^{E-09}$ | $3.8^{E-09}$ | $1.9^{E-05}$ | 0.064 |
| **N** | 121 | 110 | 53 | 99 | 106 | 104 | 59 |

### Statistical analysis of 2 h and 24 h MTZ titration (Fig. 3c)

| Compared to 0 mM MTZ | | 0.5 mM (2 h) | 1.0 mM (2 h) | 10 mM (2 h) | 0.5 mM (24 h) | 1.0 mM (24 h) | 10 mM (24 h) | Non-Tg |
|---|---|---|---|---|---|---|---|---|
| **effect size** | | -0.76 | -0.87 | -0.93 | -0.96 | -0.96 | -0.87 | -1.00 |
| **95%CI lower** | | -0.90 | -0.98 | -1.05 | -1.07 | -1.07 | -1.00 | -1.11 |
| **95%CI upper** | | -0.63 | -0.75 | -0.82 | -0.85 | -0.86 | -0.74 | -0.89 |
| ***p′*-value** | | $9.1^{E-23}$ | $1.3^{E-35}$ | $6.5^{E-41}$ | $1.5^{E-42}$ | $5.0^{E-43}$ | $9.7^{E-32}$ | $4.1^{E-45}$ |
| **N** | 232 | 110 | 149 | 129 | 77 | 119 | 72 | 330 |

| Compared to Non-Tg | | 0 mM | 0.5 mM (2 h) | 1.0 mM (2 h) | 10 mM (2 h) | 0.5 mM (24h) | 1.0 mM (24 h) | 10 mM (24 h) |
|---|---|---|---|---|---|---|---|---|
| **effect size** | | 1.00 | 0.24 | 0.13 | 0.07 | 0.04 | 0.04 | 0.13 |
| **95%CI lower** | | 0.89 | 0.15 | 0.09 | 0.04 | 0.02 | 0.02 | 0.07 |
| **95%CI upper** | | 1.11 | 0.32 | 0.18 | 0.09 | 0.06 | 0.05 | 0.20 |
| ***p′*-value** | | $4.1^{E-45}$ | $3.0^{E-06}$ | $1.5^{E-07}$ | $1.8^{E-05}$ | 0.0012 | $1.5^{E-08}$ | 0.0012 |
| **N** | 330 | 232 | 110 | 149 | 129 | 77 | 119 | 72 |

**Supplementary Table S3.** Comparisons to a signal window established by normalizing all individual sample values to the averages of the mM 0 MTZ control (set at 100%) and Non-Tg control (set at 0%) were used to calculate absolute effect sizes, 95% confidence intervals, and Bonferroni-corrected *p*-values (*p′*-values). This analysis was performed using a customized R code described in the Methods section.

#### Statistical analysis for *rho:YFP-NTR1.0* line (Fig. 4a)

| **Compared to 0 mM MTZ** | **320 nM** | **1.6 µM** | **8 µM** | **40 µm** | **0.2 mM** | **1 mM** | **5 mM** | **Non-Tg** |
|---|---|---|---|---|---|---|---|---|
| **effect size** | 0.17 | 0.02 | -0.01 | 0.23 | -0.09 | -0.70 | -0.80 | -1.00 |
| **95%CI lower** | -0.12 | -0.21 | -0.24 | 0.00 | -0.33 | -0.86 | -0.96 | -1.15 |
| **95%CI upper** | 0.46 | 0.25 | 0.22 | 0.46 | 0.14 | -0.54 | -0.65 | -0.85 |
| ***p′*-value** | 1 | 1 | 1 | 0.4 | 1 | $3.8^{E-10}$ | $1.4^{E-11}$ | $2.5^{E-13}$ |
| **N =** 29 (for 0 mM MTZ) | 29 | 28 | 28 | 28 | 28 | 28 | 27 | 38 |

| **Compared to Non-Tg** | **0 mM** | **320 nM** | **1.6 µM** | **8 µM** | **40 µm** | **0.2 mM** | **1 mM** | **5 mM** |
|---|---|---|---|---|---|---|---|---|
| **effect size** | 1.00 | 1.17 | 1.02 | 0.99 | 1.23 | 0.91 | 0.30 | 0.20 |
| **95%CI lower** | 0.85 | 0.92 | 0.83 | 0.81 | 1.04 | 0.72 | 0.23 | 0.15 |
| **95%CI upper** | 1.15 | 1.43 | 1.20 | 1.18 | 1.42 | 1.10 | 0.37 | 0.24 |
| ***p′*-value** | $2.5^{E-13}$ | $2.9^{E-09}$ | $7.4^{E-11}$ | $1.6^{E-10}$ | $1.4^{E-12}$ | $1.6^{E-09}$ | $2.1^{E-08}$ | $7.0^{E-08}$ |
| **N =** 38 (for Non-Tg) | 29 | 29 | 28 | 28 | 28 | 28 | 28 | 27 |

#### Statistical analysis for *rho:YFP-NTR2.0* line (Fig. 4b)

| **Compared to 0 mM MTZ** | **320 nM** | **1.6 µM** | **8 µM** | **40 µm** | **0.2 mM** | **1 mM** | **5 mM** | **Non-Tg** |
|---|---|---|---|---|---|---|---|---|
| **effect size** | -0.16 | -0.35 | -0.84 | -0.96 | -0.97 | -0.97 | -0.92 | -1.00 |
| **95%CI lower** | -0.35 | -0.54 | -0.98 | -1.09 | -1.10 | -1.10 | -1.05 | -1.13 |
| **95%CI upper** | 0.02 | -0.17 | -0.70 | -0.83 | -0.84 | -0.84 | -0.79 | -0.87 |
| ***p′*-value** | 0.7 | 0.002 | $4.6^{E-18}$ | $4.5^{E-20}$ | $3.6^{E-20}$ | $3.3^{E-20}$ | $2.2^{\ E-19}$ | $1.2^{E-20}$ |
| **N =** 54 (for 0 mM MTZ) | 54 | 53 | 53 | 53 | 53 | 53 | 52 | 148 |

| **Compared to Non-Tg** | **0 mM** | **320 nM** | **1.6 µM** | **8 µM** | **40 µm** | **0.2 mM** | **1 mM** | **5 mM** |
|---|---|---|---|---|---|---|---|---|
| **effect size** | 1.00 | 0.84 | 0.65 | 0.16 | 0.04 | 0.03 | 0.03 | 0.08 |
| **95%CI lower** | 0.87 | 0.70 | 0.51 | 0.11 | 0.02 | 0.01 | 0.01 | 0.05 |
| **95%CI upper** | 1.13 | 0.97 | 0.78 | 0.21 | 0.05 | 0.05 | 0.05 | 0.11 |
| ***p′*-value** | $1.2^{E-20}$ | $1.7^{E-16}$ | $1.7^{E-12}$ | $6.8^{E-07}$ | 0.002 | 0.01 | 0.02 | $1.3^{E-06}$ |
| **N =** 148 (for Non-Tg) | 54 | 54 | 53 | 53 | 53 | 53 | 53 | 52 |

**Supplementary Table S4.** Comparisons to a signal window, established by normalizing all individual sample values to the averages of the non-ablated 0 mM MTZ control (set at 100%) and Non-Tg control (set at 0%), were used to calculate absolute effect sizes, 95% confidence intervals, and Bonferroni-corrected *p′*-values. This analysis was performed using a customized R code described in the Methods section.

**# unfertilized embryos** (Fig. 5b)

| **Compared to 0 mM MTZ** | | **0.1 mM** | **1.0 mM** |
|---|---|---|---|
| **effect size** | | 2.3 | 3.2 |
| **95%CI lower** | | -2.1 | -0.8 |
| **95%CI upper** | | 6.7 | 7.3 |
| ***p*-value** | | 0.395 | 0.139 |
| **N** | 33 | 25 | 33 |

**Larval survival** (%) (Fig. 5c)

| **Compared to 0 mM MTZ** | | **0.1 mM** | **1.0 mM** |
|---|---|---|---|
| **effect size** | | 3.9% | -3.0% |
| **95%CI lower** | | -2.2% | -8.6% |
| **95%CI upper** | | 9.9% | 2.6% |
| ***p*-value** | | 0.267 | 0.387 |
| **N** | 33 | 25 | 33 |

**# fertilized embryos** (Fig. 5d)

| **Compared to 0 mM MTZ** | | **0.1 mM** | **1.0 mM** |
|---|---|---|---|
| **effect size** | | 0.2 | 6.6 |
| **95%CI lower** | | -10.1 | -2.9 |
| **95%CI upper** | | 10.5 | 16.2 |
| ***p*-value** | | 0.999 | 0.265 |
| **N** | 33 | 25 | 33 |

**Supplementary Table S5.** Statistical analysis of data presented in Fig. 5b-d. Data was subjected to ANOVA followed by Dunnett's multiple comparisons test, depending on the outcome of Levene's test. Comparisons to the 0 mM MTZ control group were used to calculate absolute effect sizes, 95% confidence intervals and *p*-values.

**Statistical analysis for *mpeg1:YFP-NTR1.1* line** (Fig. 6b)

| **Compared to 0 mM MTZ** | | **10 mM** | **1.0 mM** | **0.1 mM** |
|---|---|---|---|---|
| **effect size** | | -0.29 | -0.05 | 0.02 |
| **95%CI lower** | | -0.41 | -0.16 | -0.07 |
| **95%CI upper** | | -0.18 | 0.07 | 0.11 |
| ***p′*-value** | | $2.7^{E-04}$ | 1 | 1 |
| **N** | 9 | 9 | 8 | 7 |

**Statistical analysis for *mpeg:Gal4;UAS:YFP-NTR2.0*** (Fig. 6d)

| **Compared to 0 mM MTZ** | | **10 mM** | **1.0 mM** | **0.1 mM** |
|---|---|---|---|---|
| **effect size** | | -0.64 | -0.73 | -0.67 |
| **95%CI lower** | | -0.85 | -0.85 | -0.85 |
| **95%CI upper** | | -0.43 | -0.62 | -0.49 |
| ***p′*-value** | | $7.2^{E-05}$ | $9.2^{E-09}$ | $4.4^{E-06}$ |
| **N** | 10 | 9 | 9 | 9 |

**Supplementary Table S6:** Comparisons to 0 mM MTZ controls were used to calculate absolute effect sizes, 95% confidence intervals, and Bonferroni-corrected *p′*-values. This analysis was performed using a customized R code described in the Methods section.

Supplementary Fig. S5: NTR 2.0 does not enhance ablation efficacy with nifurpirinol.

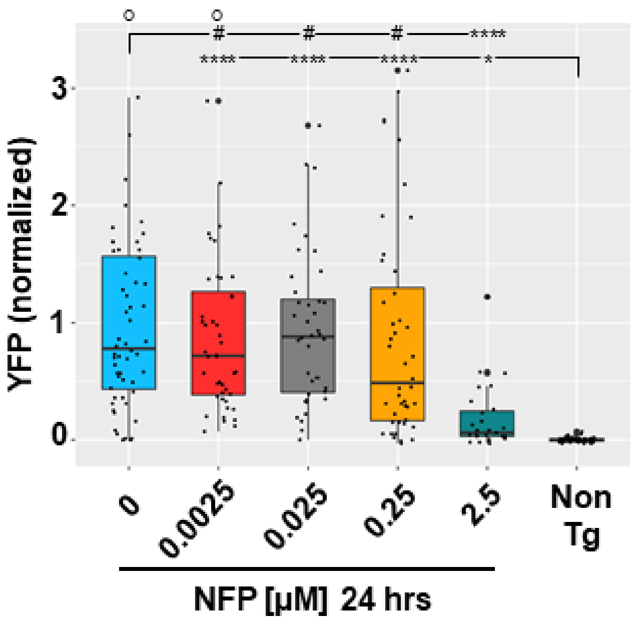


Double transgenic zebrafish larvae co-expressing YFP and NTR 2.0 were exposed to the indicated concentrations of NFP from 5 to 6 dpf and YFP levels quantified by plate reader at 7 dpf. Statistical comparisons to two controls, non-ablated (0 mM MTZ) and age-matched non-transgenic (Non-Tg) larvae, were used to calculate effects sizes, 95% confidence intervals, and Bonferroni-corrected *p'*-values (provided in Supplementary Table S7; $^{\#}p>0.05$, $^{*}p<0.05$, **** $p<0.0001$; ○ = data points beyond the bounds of the graph).

**Statistical analysis for 24 h NFP** (Fig. S5)

| Compared to 0 µM NFP | | 0.0025 µM | 0.025 µM | 0.25 µM | 2.5 µM | Non-Tg |
|---|---|---|---|---|---|---|
| **effect size** | | -0.09 | -0.06 | -0.12 | -0.82 | -1.00 |
| **95%CI lower** | | -0.39 | -0.35 | -0.47 | -1.05 | -1.21 |
| **95%CI upper** | | 0.22 | 0.23 | 0.22 | -0.58 | -0.79 |
| ***p′*-value** | | 1 | 1 | 1 | $6.3^{E-09}$ | $1.9^{E-12}$ |
| **N** | 56 | 44 | 40 | 44 | 27 | 83 |
| **Compared to Non-Tg** | | **0 µM** | **0.0025 µM** | **0.025 µM** | **0.25 µM** | **2.5 µM** |
| **effect size** | | 1.00 | 0.91 | 0.94 | 0.88 | 0.18 |
| **95%CI lower** | | 0.79 | 0.69 | 0.73 | 0.60 | 0.07 |
| **95%CI upper** | | 1.21 | 1.14 | 1.15 | 1.15 | 0.29 |
| ***p′*-value** | | $1.9^{E-12}$ | $1.5^{E-09}$ | $1.5^{E-10}$ | $4.8^{E-07}$ | 0.011 |
| **N** | 83 | 56 | 44 | 40 | 44 | 27 |

**Supplementary Table S7**: Comparisons to a signal window established by normalizing all individual sample values to the averages of the 0 µM NFP controls (set at 100%) and Non-Tg controls (set at 0%) were used to calculate absolute effect sizes, 95% confidence intervals, and Bonferroni-corrected *p′*-values. This analysis was performed using a customized R code described in the Methods section.

| Construct/Transgenic line and allele | Expression system | NTR co-expressed reporter | "Tracer" reporter |
|---|---|---|---|
| *Tg(5xUAS:GAP-EGFP-P2A-NfsB_Vv F70A/F108Y,he:tagBFP2)jh551* | Gal4/UAS | EGFP | tagBFP2 |
| *Tg(5xUAS:GAP-tagYFP-P2A-NfsB_Vv F70A/F108Y,he:tagBFP2)jh552* | Gal4/UAS | tagYFP | tagBFP2 |
| *Tg(5xUAS:GAP-mCherry-P2A-NfsB_Vv F70A/F108Y,he:tagBFP2)jh553* | Gal4/UAS | mCherry | tagBFP2 |
| *Tg(5xQUAS:GAP-tagYFP-P2A-NfsB_Vv F70A/F108Y,he:ECFP)jh562* | QF/QUAS | YFP | ECFP |

**Supplementary Table S8.** List of bipartite reporter/effector plasmids and transgenic resources for driver-defined expression of NTR 2.0.

| Primer Name | Sequence (5' → 3') |
|---|---|
| ***Primers for insertion of nitroreductase genes into vectors pUCX and pET28a+*** | |
| NfsB_Vv Fw | GGGGCATATGACTATTGTTCAAGCT |
| NfsB_Vv Rv | GGGGTCGACTTAGATTTCGGTAAAAACAG |
| NfsB_Ec Fw | GGGCATATGGATATCATTTCTG |
| NfsB_Ec Rv | GGGGAATTCTTACACTTCGGTTAAG |
| NfsB_Kp_Fw | GGGGCATATGGATATCGTATCGGTC |
| NfsB_Kp_Rv | GGGGCTCGAGTTAAATCTCGGTAATGGT |
| NfsB_Pp_Fw | GGGGCATATGGATACCGTATCGCTG |
| NfsB_Pp_Rv | GGGGCTCGAGTCAGAGGAAGGTGAACACTT |
| NfsB_St_Fw | GGGGCATATGGATATCGTTTCTGTC |
| NfsB_St_Rv | GGGGGTCGACTTAGACTTCCGTCAGTG |
| NfsB_Ck_Fw | CCCCCATATGGATATCGTTTCTGTCGC |
| NfsB_Ck_Rv | CCCGTCGACTCAGACTTCCGTCAGGGTG |
| FraseI_Vf Fw | GGGGCATATGACGCATCCAATTATTCA |
| FraseI_Vf Rv | GGGGGTCGACCTAAAGAATGGTAATTAC |
| NfsB_Vh_Fw | CCCCCATATGTCTCATCAAATCATTACAGAC |
| NfsB_Vh_Rv | CCCCTCGAGTTAAAGAGTAGTAATTACGTC |
| YfkO_Bs_Fw | GGGGCATATGGCAGATCTAAAGACACA |
| YfkO_Bs_Rv | CTAGGTCGACTTAAACCCACTTCACAACAT |
| YdgI_Bs_Fw | GGGGCATATGATCAAAACAAACGATTTTATGG |
| YdgI_Bs_Rv | GGGGGTCGACTTATTTCCATTCTGCAATTG |
| NfsB_Cs_Fw | CCCCCATATGAACCTTAATGAGATCATTCGC |
| NfsB_Cs_Rv | CCCGTCGACTCAGAGCTGCGTGATCAC |
| ***Primers for insertion of nitroreductase genes into vector F279-V5*** | |
| NfsB_Vv GW_Fw | GGGGACAAGTTTGTACAAAAAAGCAGGCTTCGAAGGAGATAGAACCATGG GCACTATTGTTCAAGCT |
| NfsB_Vv GW_Rv | GGGGACCACTTTGTACAAGAAAGCTGGGTCCTAGATTTCGGTAAA |
| NfsB_Ec GW_Fw | GGGGACAAGTTTGTACAAAAAAGCAGGCTTCGAAGGAGATAGAACCATGG GCGATATCATTTCTGTCGCCTT |
| NfsB_Ec GW_Rv | GGGGACCACTTTGTACAAGAAAGCTGGGTCCTACACTTCGGTTAAGGTGA TGT |
| ***Primers for insertion of nitroreductase variants into vector pTOL2-5xUAS-GAP-tagYFP-P2A-nfsB_Ec*** | |
| NfsB_Vv pTOL2_Fw | CCCCCCGGGCCTATGACTATTGTTCAA |
| NfsB_Vv_Rv | GGGGTCGACTTAGATTTCGGTAAAAACAG |
| ***Primers for insertion of rhodopsin promoter into vector pTOL2-5xUAS-GAP-tagYFP-P2A-nfsB_Ec*** | |
| rho_promoter_F2 | AATTTAGGGTAATGGCCGGCCCGACCTGCAGGTCAACGG |
| rho_promoter_R | AGTCTACCCTGCTACGTAGGCTGCGGTTGGATGTGG |
| ***Primers for swapping eGFP gene into 5xUAS:YFP-2A-NTR2.0 vector*** | |
| SbfI-GFP_FOR | GCAGCCTGCAGGAAGCACAGCTACAATGATGGTGAGCAAGGGCGAG |
| GFP-oe_REV | CTCCGCTTCCCTTGTACAAGTACTTGTACAGCTCGTCCATGC |
| FW-P2A | TACTTGTACAAGGGAAGCGGAGCTA |
| KflI-Bglobin_REV | CAAGGGTCCCCAAACTCACC |
| ***Primers for insertion of YFP-2A-NTR 2.0 gene into 5xQUAS vector*** | |
| mluI-YFP_FOR | TTAACGCGTAGCAGGGTAGACTGTAGAGG |
| AvrII-He1Pro_REV | GGCCCTAGGGTTGCCTCA |

**Supplementary Table S9.** Primers used in this study. Underlined sequences are restriction, recombination, or overlap extension sites.

| **Individual Transgenic Lines** | | | |
|---|---|---|---|
| **Full Transgene Name** | **NTR** | **Abbreviation** (here) | **Ref.** |
| Tg(5xUAS:GAP-tagYFP-P2A-nfsB_Vv F70A/F108Y)jh513 | 2.0 | UAS:YFP-NTR2.0 | here |
| Et(2xNRSE-fos:KALTA4)gmc617 | na | NRSE:KalTA4 | [26] |
| Tg(5xUAS:GAP-ECFP,he:GAP-ECFP)gmc1913 | na | UAS:CFP | here |
| Tg(14xUAS:nfsB_Ec-mCherry)c264 | 1.0 | UAS:NTR1.0-Cherry | [28] |
| Tg(mpeg1.1:EYFP-nfsB_Ec T41Q/N71S/F124T)w202 | 1.1 | mpeg:NTR1.1-YFP | [20] |
| Tg(rho:EYFP-nfsB_Ec)gmc500 | 1.0 | rho:YFP-NTR1.0 | [15] |
| Tg(rho:GAP-tagYFP-P2A-nfsB_Vv F70A/F108Y)jh405 | 2.0 | rho:YFP-NTR2.0 | here |
| Tg(mpeg1.1:Gal4-VP16)gl24 | na | mpeg:Gal4 | [49] |
| **Double Transgenic Lines** | | | |
| **Full Transgene Name** | **NTR** | **Abbreviation** (here) | Ref. |
| Et(2xNRSE-fos:KALTA4)gmc617;<br>Tg(5xUAS:GAP-tagYFP-P2A-nfsB_Vv F70A/F108Y)jh513 | 2.0 | NRSE:KalTA4;<br>UAS:YFP-NTR2.0 | [26]<br>here |
| Et(2xNRSE-fos:KALTA4)gmc617;<br>Tg(5xUAS:GAP-ECFP,he:GAP-ECFP)gmc1913 | na | NRSE:KalTA4;<br>UAS:CFP | [26]<br>here |
| Tg(mpeg1.1:Gal4-VP16)gl24;<br>Tg(5xUAS:GAP-tagYFP-P2A-nfsB_Vv F70A/F108Y)jh513 | 2.0 | mpeg:Gal4;<br>UAS:YFP-NTR2.0 | [49]<br>here |
| **Triple Transgenic Lines** | | | |
| **Full Transgene Name** | **NTR** | **Abbreviation** (here) | Ref. |
| Et(2xNRSE-fos:KALTA4)gmc617;<br>Tg(5xUAS:GAP-tagYFP-P2A-nfsB_Vv F70A/F108Y)jh513;<br>Tg(5xUAS:GAP-ECFP,he:GAP-ECFP)gmc1913 | 2.0 | NRSE:KalTA4;<br>UAS:YFP-NTR2.0<br>UAS:CFP | [26]<br>here<br>here |
| Et(2xNRSE-fos:KALTA4)gmc617;<br>Tg(5xUAS:GAP-tagYFP-P2A-nfsB_Vv F70A/F108Y)jh513;<br>Tg(14xUAS:nfsB_Ec-mCherry)c264 | 2.0<br>1.0 | NRSE:KalTA4;<br>UAS:YFP-NTR2.0<br>UAS:NTR1.0-Cherry | [26]<br>here<br>[28] |
| **New Reporter/Effector Transgenic Lines** | | | |
| **Full Transgene Name** | **NTR** | **Abbreviation** (here) | **Ref.** |
| Tg(5xUAS:GAP-EGFP-P2A-NfsB_Vv F70A/F108Y,he:tagBFP2)jh551 | 2.0 | na | here |
| Tg(5xUAS:GAP-tagYFP-P2A-NfsB_Vv F70A/F108Y,he:tagBFP2)jh552 | 2.0 | na | here |
| Tg(5xUAS:GAP-mCherry-P2A-NfsB_Vv F70A/F108Y,he:tagBFP2)jh553 | 2.0 | na | here |
| Tg(5xQUAS:GAP-tagYFP-P2A-NfsB_Vv F70A/F108Y,he:ECFP)jh562 | 2.0 | na | here |

**Supplementary Table S10.** Transgenic lines used in this study.
